# Enhanced Radio-sensitization of Glioblastoma using a Dendrimer-Based Metformin Nano-formulation through Direct Tumor Suppression and Indirect Immune Modulation

**DOI:** 10.64898/2026.05.12.724405

**Authors:** Sadaf Mahfooz, Fei Wang, Ghanbar Mahmoodi Chalbatani, Tatiana K Bronich, Svetlana Romanova, Yuanyuan Jia, Krishna Bhat, Chi Zhang

## Abstract

Glioblastoma (GBM) is the most common and lethal primary malignant brain tumor in adults, with median survival remaining approximately 12–15 months despite aggressive multimodal therapy. Therapeutic resistance and tumor recurrence are driven in part by limited drug penetration across the blood–brain barrier (BBB) and the persistence of brain cancer stem cells (BCSCs), highlighting the need for brain-penetrant therapeutic platforms capable of achieving sustained intratumoral delivery. Here, we developed a dendrimer-based nanotherapeutic by conjugating metformin to a fourth-generation hydroxyl-terminated polyamidoamine dendrimer (P4-MET) to enhance intracranial bioavailability and therapeutic efficacy in GBM.

P4-MET exhibited favorable pharmacokinetic properties, including prolonged retention within the tumor microenvironment, and demonstrated enhanced cytotoxicity against GBM cell lines relative to free metformin (f-MET). Mechanistical studies with transcriptomic profiling by RNA sequencing revealed distinct treatment-associated molecular signatures, identifying BOLA2B as the most significantly differentially expressed gene between treatment groups. Specifically, BOLA2B expression was markedly elevated in f-MET–treated cells but not so following P4-MET treatment. Given the established association of BOLA2B with mTORC1 signaling and GPX4-mediated ferroptosis resistance, these findings suggest that P4-MET may, at least in part, enhance therapeutic efficacy by modulating ferroptosis-associated pathways.

In orthotopic GBM models, combination treatment with P4-MET and radiotherapy (RT) significantly prolonged overall survival and increased tumor cell death compared with either monotherapy alone, consistent with a synergistic radiosensitizing effect. Importantly, P4-MET demonstrated minimal systemic toxicity, supporting its favorable therapeutic index and translational potential. Collectively, these findings establish P4-MET as a brain-penetrant nanomedicine platform that improves metformin delivery, modulates ferroptosis-related signaling networks, and potentiates radiotherapeutic response in GBM. This study highlights the potential of dendrimer-enabled metabolic nanotherapies to overcome therapeutic resistance in malignant brain tumors.

## Introduction

For more than two decades, the standard of care for glioblastoma (GBM), the most aggressive primary brain tumor in adults, has remained essentially unchanged. The treatment typically begins with maximal safe surgical resection, followed by adjuvant radiotherapy and chemotherapy with temozolomide (TMZ), an alkylating agent that disrupts tumor cell replication by inducing DNA damage through base mismatches and strand breaks. This drug is particularly effective during oncogenesis, where rapid cellular proliferation renders cancer cells more vulnerable to DNA-targeting agents. Radiotherapy acts synergistically with TMZ by causing additional DNA damage, leading to increased tumor cell death. Although it can cross the BBB to exert its therapeutic effects, achieving effective concentration often requires high systemic doses, which are associated with significant adverse effects, including myelosuppression, gastrointestinal toxicity, and fatigue ^1, 2^.

Furthermore, despite this aggressive multimodal approach, the prognosis for GBM patients remains dismal, with a median survival of only 16 to 18 months. The primary reason for poor outcomes is tumor recurrence, driven by residual infiltrative tumor cells and the presence of therapy-resistant brain cancer stem cells. These limitations underscore the urgent need for innovative therapeutic strategies to enhance treatment efficacy while minimizing systemic toxicity ^3^_._

Metformin, a widely used antidiabetic drug, has demonstrated antitumor properties in various cancers, including GBM, and in immune cells, through mechanisms such as activation of AMP-activated protein kinase (AMPK), inhibition of mitochondrial complex I, reduction of cellular proliferation, and modulation of cancer stem cell signaling pathways ^4–6^. The activated AMPK further inhibits the mammalian target of rapamycin (mTOR) and its downstream effector molecules, thereby enhancing tumor cell death^7^. The antitumor effect of metformin has been reported in various in vitro and in vivo studies, highlighting AMPK activation as its critical mechanism of action^8, 9^.

Advances in nanotechnology, targeted drug delivery systems, and novel drug formulations are now being explored to address these challenges. By improving BBB penetration, selectively targeting tumor cells, and enhancing drug accumulation within the tumor microenvironment, these approaches have the potential to significantly improve outcomes for GBM patients^10, 11^. Polyamidoamine (PAMAM) dendrimers have garnered substantial attention for their versatility and reproducibility as nanocarriers that can be modified and loaded with drugs or genes to target cancer cells^12^. Recently, numerous studies have employed PAMAM dendrimer-based nanoformulations to encapsulate drugs and target tumor cells, including GBM, as an alternative therapeutic approach^13, 14^.

We developed and characterized a novel nano-formulation (P4-MET) consisting of fourth-generation polyamidoamine [PAMAM-OH (G4)] dendrimers conjugated to metformin via a stable carbamate bond. This conjugation strategy enables controlled intracellular release of metformin in response to acidic and enzymatic conditions. The PAMAM dendrimer platform offers enhanced biocompatibility, efficient penetration across the BBB, and prolonged tumor retention, thereby optimizing metformin pharmacokinetics. We evaluated the therapeutic potential of P4-MET in the CT-2Aluc murine glioblastoma model, assessing its ability to inhibit tumor growth, modulate metabolic pathways, and improve survival. Our findings provide critical insights into the potential of dendrimer-based nano-formulations as an innovative strategy for GBM treatment. For the first time, the nano-formulation of metformin (P4-MET) has been used in the present study to target GBM.

## Materials and Methods

### Materials

All reactions in the organic medium were performed in standard oven-dried glassware under an inert nitrogen atmosphere using freshly distilled solvents. PAMAM-OH dendrimer (biomedical-grade generation 4 consisting of 64 hydroxyl end-groups) (G4-OH) was purchased from Sigma-Aldrich. Spectra/Por Dialysis membrane (MWCO: 3,500–5,000 Da) was purchased from Spectrum Laboratories, a division of Thomas Scientific. The dendrimers were purified before use. IR Dye 800CW NHS Ester (IR800-NHS) was obtained from LI-COR Biosciences, and Alexa Fluor 488 NHS Ester (A488-NHS) was purchased from Thermo Fisher Scientific. All other reagents were obtained from Sigma-Aldrich and used as supplied without prior purification unless otherwise stated. For biological studies, fetal bovine serum (FBS), Dulbecco’s Modified Eagle Medium (DMEM), sodium hydroxide (NaOH), metformin hydrochloride (f-MET), trypsin–EDTA solution (0.25%), acetonitrile (ACN), dimethyl sulfoxide (DMSO), and methanol were purchased from Sigma-Aldrich. BOLA2B (Gene ID-654483) SMARTpool siRNAs were procured from Dharmacon. Deferoxamine mesylate [DFO] (Cat#sc-203331), a BOLA2B inhibitor, was purchased from Santa Cruz Biotechnology (SCBT).

### Synthesis of Free Base Metformin

Metformin hydrochloride (20 g, 0.12 mol) was dissolved in 1N NaOH (120 mL, 0.12 mol). The reaction mixture was stirred at room temperature (RT) for 30 minutes, then concentrated under reduced pressure. Ethanol (50 mL) was added, and the mixture was stirred for 30 minutes. The solution was then filtered and concentrated under a vacuum. This procedure was repeated twice to ensure the complete removal of residual salt.

### Synthesis of P4-MET Conjugate

Methanol was removed under reduced pressure, yielding a viscous oil of PAMAM-OH dendrimer. Dendrimer (3.25 g, 0.01464 mmol of hydroxyl groups) was dissolved in a 10 mL mixture of anhydrous dimethyl sulfoxide (DMSO) and dimethylformamide (DMF) mixture (3:1, v/v). Hydroxyl groups were activated using carbonyl diimidazole (CDI) (1.5 equivalents per hydroxyl group). CDI (3.56 g, 0.02196 mmol) was dissolved in 5 mL of anhydrous DMSO, and the dendrimer solution was added dropwise under argon, followed by stirring at RT for 24 hours. Free-base metformin (8 equivalents per activated hydroxyl group) was then introduced, and the reaction continued under argon for 48 hours. The final product was purified via dialysis against distilled water for 48 hours, followed by an additional 24-hour dialysis against phosphate-buffered saline (PBS, pH 7.4). The purified conjugate was then freeze-dried and further purified by flash column chromatography on LH-20 resin using methanol as the eluent.

### Synthesis of fluorescently-labeled-conjugates

P4-MET conjugate (0.8 g, 0.0476 mmol; ∼3 mmol of hydroxyl groups) was dissolved in anhydrous DMSO and reacted with Boc-amino-undecanoic acid to yield PAMAM-MET-DEC-NH-Boc. The product was purified by dialysis against DMSO (molecular weight cutoff: 1000 Da), followed by dialysis against water. After drying, the Boc-protecting group was removed using trifluoroacetic acid in dichloromethane. The resultant PAMAM-MET-DEC-NH_2_ dendrimer was isolated via co-evaporation with dichloromethane (three cycles). The purified amine-terminated dendrimer was labeled with A488 or IR800 and purified via size-exclusion chromatography (LH20; methanol).

### 1H-NMR Spectroscopy

Proton nuclear magnetic resonance (^1^H-NMR) spectroscopy confirmed the chemical structure and purity of the intermediates (P4-CDI) and the final P4-MET conjugate. Spectra were recorded at 400 MHz using a high-resolution NMR spectrometer, with deuterated dimethyl sulfoxide (DMSO-d6) and D_2_O as the solvents for the P4-CDI and P4-MET, respectively. Characteristic peaks corresponding to methyl (–CH3), methylene (–CH2), and amide (–NH) groups of PAMAM and metformin were identified. Chemical shifts associated with the dendrimer and metformin confirmed successful conjugation via carbamate linkage. Peak integration was further analyzed to estimate the metformin grafting density on the PAMAM scaffold.

### Dynamic Light Scattering (DLS) and Zeta Potential Analysis

The hydrodynamic diameter (d, nm) and polydispersity index (PDI) of P4-MET nanoparticles were measured using dynamic light scattering (DLS) in PBS at 25°C with a fixed scattering angle of 173° (Nano ZS Zetasizer, Malvern Instruments, UK). Zeta potential measurements were conducted to assess colloidal stability. Data was acquired using Zetasizer software (Version 7.11) and reported as the means of three independent measurements.

### Transmission Electron Microscopy (TEM)

Nanoparticle morphology was analyzed using transmission electron microscopy (TEM). Samples for TEM imaging were spotted onto formvar/silicon monoxide-coated 200 mesh copper grids (Ted Pella Inc., Redding, CA). Grids were glow-discharged for 60 seconds at 20µA with a GloQube glow discharge unit (Quorum Technologies, East Sussex, UK) before use. Samples were negatively stained with NanoVan (Nanoprobes, New York, NY) and examined on a Tecnai G2 Spirit TWIN (FEI, Hillsboro, OR) operated at 80kV. Images were acquired digitally with an AMT (Woburn, MA) digital imaging system.

The biodistribution of P4-MET in the CT-2Aluc cells was also evaluated by TEM imaging. For this purpose, the cells were fixed by immersion in a solution of 2% glutaraldehyde, 2% paraformaldehyde in a 0.1M Sorenson’s phosphate buffer (pH 6.2) for a minimum of 24 h at 4°C. Samples were then washed three times with PBS to clear excess fixative. During processing, samples were post-fixed in a 1% aqueous solution of osmium tetroxide for 30 minutes. Subsequently, the samples were dehydrated in a graded ethanol series (50%, 70%, 90%, 95%, 100%), with propylene oxide used as a transition solvent between ethanol and Araldite resin. Samples were allowed to sit overnight in a 50:50 propylene oxide: resin solution until all propylene oxide had evaporated. Samples were then incubated in fresh resin for 2 hours at room temperature before final embedding. Polymerization took place at 65°C for 24 hours. Thin sections (100nm) prepared with a Leica UC6 Ultra cut ultramicrotome were placed on 200-mesh copper grids and examined using a Tecnai G^2^ Spirit TWIN (FEI) operating at 80kV. No additional staining was performed for this experiment. Images were acquired digitally with an AMT digital imaging system.

### Encapsulation Efficiency and Drug Loading Analysis

The encapsulation efficiency (EE) and drug loading capacity of P4-MET were quantified using ultra-performance liquid chromatography (UPLC). Analyses were performed on a Waters ACQUITY UPLC system equipped with a BEH C18 column (Waters ACQUITY UPLC® BEH C18, 1.7 µm, 2.1 × 50 mm). The mobile phase consisted of solvent A (water with 0.1% formic acid) and solvent B (acetonitrile with 0.1% formic acid), using a gradient elution method. The flow rate was maintained at 0.3 mL/min, with a column temperature of 40°C and an injection volume of 1 μL. Detection was performed using a photodiode array (PDA) detector at 230 nm. Samples were filtered through a 0.22 µm membrane before injection.

Encapsulation efficiency (EE%) was calculated as:

EE (%) = (Amount of drug encapsulated/Total amount of drug added) x 100

Drug loading capacity (DL%) was calculated as:

DL (%) = (Amount of drug encapsulated/Total weight of nanoparticles) x 100

Calibration curves for free metformin were generated using known concentrations (0-100μg/ml), and sample concentrations were quantified using the curve’s linear regression equation.

### Metformin Release Studies

A release study was conducted to evaluate the drug release profile of the P4-MET conjugate under various physiological conditions, simulating both intracellular and extracellular environments. The conjugate was placed in dialysis bags and immersed in PBS at pH 7.4 (physiological conditions), pH 5.5, and at a low pH (4.5) (mimicking the tumor microenvironment). Samples were collected at predetermined intervals and analyzed by UPLC to quantify metformin.

### Endotoxin Limit Calculation

The endotoxin limit calculations were performed by the Nanotechnology Characterization Laboratory (NCL) in the USA, based on several assumptions, including that the entire injected dose remains in circulation, for a 70 kg patient. If the dose changes, Human Equivalent dose (HED), Theoretical Human Plasma Concentration at HED (THPC), and Endotoxin Limit (EL) will also change.

The in vitro test concentrations used were 5, 0.5, 0.1, and 0.02 mg/mL of the API (metformin).

HED = mouse dose ÷ 12.3

THPC = HED x 70 kg ÷ 5.6 L

EL = K/M; where K = 5 EU/kg, M = HED

MVD = (EL x sample concentration)/λ

[λ = 0.001 EU/mL for turbidity and chromogenic and 0.03 EU/mL for gel clot]

### Cell Culture

U87MG (TP53 wildtype and PTEN mutant), Ln229, SF188, and CT-2Aluc were purchased from the American Type Culture Collection (ATCC) (Manassas, VA). A murine cell line MGPP3 (PDGF+, P53-/-, PTEN -/-) was kindly provided by Dr. Peter Canoll (Columbia University, New York, NY). Primary GBM cells were isolated from an adult patient’s GBM tumor sample. Human GBM cells were cultured in DMEM (ATCC) according to the provider’s recommendations. Murine GBM cells were cultured using the previously published protocol ^15^. All experiments were performed with cells in passages 3-6. Cell culture experiments were conducted under standard sterile conditions at 37°C in a humidified 5% CO₂ incubator.

### Cell Viability assay

The Cell Titer Blue (CTB) assay was used to evaluate the viability of GBM cells cultured in 96-well plates and treated with free metformin (f-MET), P4-OH, PAMAM mixed with free metformin (P4-OH+f-MET), and P4-MET at concentrations equivalent to the free metformin content for 72 hours. After 72 hours, 20 µL of Cell Titer Blue reagent (Promega, Madison, WI, USA) was added to each well, replacing the drug-containing medium with 100 µL of fresh medium. The mixture was then incubated for 2 hours at 37°C. After incubation, fluorescence was recorded using a SpectraMax (Molecular Devices, San Jose, CA, USA) at excitation and emission wavelengths of 560 nm and 590 nm, respectively. IC_50_ values for each treatment group in different GBM cell lines were calculated through GraphPad software (Version 8.3.1, San Diego, CA, USA).

### Cellular uptake study by confocal microscopy

To further evaluate the in vitro cellular uptake of the drug, CT-2Aluc cells were plated in 12-well plates with coverslips and cultured overnight. Afterward, CT-2Aluc cells were treated with the fluorescently labeled drug P4-MET-AF488 for 4h, 16h, and 24h. After drug treatment, remove the medium and wash the cells twice with 1X PBS. Cells were then labeled with Mitotracker by adding 200nM Mitotracker to the medium and incubating for 30 minutes. After incubation, the cells were mounted in VECTASHIELD containing 4′,6-diamidino-2-phenylindole (DAPI). Images demonstrating cellular uptake were captured at 63× magnification using a confocal laser scanning microscope (Carl Zeiss, LSM 710/800 META).

### Immunoblotting

The CT-2Aluc cells were treated with 1 mM free metformin and P4-MET at equivalent concentrations for 4, 8, 24, and 48 hours. After treatment, the cell pellets were homogenized using RIPA Lysis and Extraction Buffer (Thermo Scientific Inc., Waltham, MA, USA) combined with a protease inhibitor cocktail (Thermo Fisher Scientific Inc., USA). The protein concentrations were quantified using the Pierce BCA Protein Assay kit (Thermo Scientific) according to the manufacturer’s protocol, and fluorescence was measured with a spectrophotometer (Infinite 200Pro). The cell lysates, containing 30-40µg of protein, were separated using 4-20% Tris-glycine SDS-PAGE gels (Thermo Fisher Scientific Inc.). The primary antibodies used in the present study include anti-phospho-AMPKa (Thr172) (Cat # 2535s), Recombinant Anti-AMPK alpha 1 + AMPK alpha 2 antibody (cat # ab207442), AKT Monoclonal Antibody (2C5D1), Phospho-Akt (Ser473) (587F11) Mouse mAB, Phospho-Akt (Thr308) (D25E6) XP® Rabbit mAb, Phospho-mTOR (Ser2448) (D9C2) XP® Rabbit mAb (Cat#5536s), Recombinant Anti-mTOR antibody (Cat#ab32028), S6 Ribosomal (Cat#14798), pS6Ribosomal Ser235/236 (Cat#4858S), Cyclin D1 mouse monoclonal (DCS-6) (Cat#SC-20044), REDD1 rabbit polyclonal (Cat#10638-1-ap), BOLA2 rabbit polyclonalAb (Cat#26080-I-AP), and anti-β-actin (AC-74, 1:3000) (Millipore Sigma). After incubating the PVDF membrane with the primary antibodies overnight at 4°C, the membranes were washed three times with TBST for 5 minutes each. After washing, the membranes were incubated with the IRDye 800CW Goat Anti-Mouse IgG (H+L) or IRDye 800CW Goat Anti-rabbit IgG (H+L) secondary antibodies (Li-Cor biotechnology, Lincoln, NE, USA) at room temperature for 1h. The images of the blots were captured using the ChemiDoc Imaging System (BIO-RAD). ImageJ software was used to quantify protein expression relative to the control.

### Quantitative Real-Time (qRT-PCR)

Total RNA from GBM22 cells was isolated using the RNeasy Kit (Qiagen, MD, USA), followed by NanoDrop quantification and cDNA synthesis with the iScript cDNA synthesis kit (Bio-Rad, USA), as previously described. Gene expression of BOLA2b was evaluated using Fast SYBR Green Master Mix (Applied Biosystems, USA) by quantitative real-time PCR (qRT-PCR) on a QuantStudio 6 Flex (Applied Biosystems, USA) following the manufacturer’s protocol. Gene expression was normalized to the RPL13A internal control. The primer sequences used are mentioned below, For hBOLA2B: F-TTCAGAGACACAGGCTGGTGA; R- GGGGTCAGGGTTTTCTGTTCA For hRPL13A: F-CTAAACAGGTACTGCTGGGC; R- AGGAAAGCCAGGTACTTCAACTT

### Seahorse ATP production rate assay

Glycolytic and mitochondrial ATP production rates were quantified after the metformin and P4-MET treatment in CT-2Aluc cells by the Seahorse XFe96/XF Pro assay (Agilent) following the manufacturer’s protocol. Briefly, the 15×10^3^cells were seeded in the specialized Seahorse 96-well plate. Thereafter, prepare the seahorse sensor cartridge and replace it with the seahorse calibrant. Then, the cells were treated with 1 mM f-MET and an equivalent dose of P4-MET for 48 h. After treatment, extracellular acidification rate (ECAR) and oxygen consumption rate (OCR) were measured under basal settings. Then, the oligomycin and rotenone/antimycin A were consecutively added to distinguish the contribution of ATP through oxidative phosphorylation and glycolysis, respectively.

### Syngeneic orthotopic GBM mouse model development

Orthotopic syngeneic mouse GBM models have been established using a stereotactic microinjection technique routinely performed in our laboratory ^17^. We implanted 5 × 10^^4^ CT-2Aluc cells (Millipore-Sigma), which are stably transfected with a luciferase reporter gene and are a model cell line for GBM studies, into the right brain hemisphere of healthy female Albino B6 mice, approximately 12 weeks old and weighing 20-30 g. Tumor growth was confirmed by brain imaging using the LI-COR Pearl Trilogy imaging system at 10 min after a 150 mg/kg IP luciferin injection, weekly. P4-MET or f-MET (200mg/kg/d, equivalent to free metformin) was administered daily by Intraperitoneal (IP) injection starting on day 8 post-implantation for 3 weeks, with or without whole brain RT, to a total dose of 8Gy in one fraction on day 11. Overall survival (OS) across groups (8-12 mice per group) was compared. After tagging P4-MET with the infrared fluorescent dye IR800 (Li-Cor) for injection and live whole-body fluorescent scanning using the Pearl Trilogy Imaging System (Li-Cor, Lincoln, NE), we also investigated the pharmacodynamics of P4-MET penetration into tumor-bearing tissue (in the brain). Immunohistochemistry (IHC) was performed by perfusing the dead mice with PBS containing 10% formalin, followed by dissection of the brain tissues for staining of infiltrating immune cell markers after P4-MET treatment. All experiments were conducted in accordance with the in-house ethical guidelines for animal studies. All the animals received medications to control pain after the surgery, throughout the follow-up, and before euthanization according to the guidelines of the institute. The protocol was approved by the Institutional Animal Care and Use Committee (IACUC) of the University of Nebraska Medical Center (UNMC).

### Survival study

Survival analysis was conducted to evaluate the therapeutic efficacy of f-MET and P4-MET in a mouse model of glioblastoma. The primary objective was to assess whether treatment with these formulations could prolong survival compared to control groups, particularly in the context of glioblastoma progression and the potential synergy with radiotherapy. Mice-bearing established GBM tumors were randomly assigned to different treatment groups: (1) Control (Normal saline), (2) f-MET, (3) P4-MET, (4) radiotherapy (RT) alone, (5) combination of f-MET and RT, and (6) combination of P4-MET and RT. The treatments were administered over a defined period, with survival monitored as the primary endpoint. Mice were observed daily for signs of distress, weight loss, or other clinical symptoms of tumor progression, and survival was tracked until humane endpoints were reached. The Kaplan-Meier method was used to estimate overall survival in each treatment group, and statistical analysis was performed to compare survival curves.

### Pharmacokinetics (PK) and biodistribution study

The pharmacokinetics of P4-MET were determined by measuring the fluorescent signal in tissue at various time points after IP injection of fluorescently labeled formulations in mice. Healthy mice (weight: 20 ± 2 g, n = 5) were treated with P4-MET at a concentration of 200mg/kg. Additionally, metformin concentrations in blood plasma, liver, and brain tissue were analyzed using a mass spectrometer (Agilent 1290 UPLC-6495 QQQ MS). Biodistribution of P4-MET in organs/tissues was evaluated via fluorescence imaging of these organs/tissues after injection of fluorescently labeled formulations into mice bearing a GBM tumor. The mice with a tumor were randomly divided into two groups. These mice were injected with P4-MET at a dose of 200 mg/kg through the IP route. Three mice of each group were continuously imaged using fluorescence at 1, 6, 12, 24, 48, and 72 h post-injection. To further characterize the distribution of nano-formulations, three mice from each group were simultaneously dissected after injection. The major organs (brain, heart, liver, spleen, lung, and kidney) were collected and weighed. Semi-quantitative analysis of fluorescence images of these organs/tissues was performed using the Pearl Trilogy imaging system. Mice injected with normal saline served as the control. The tissue penetration of P4-MET was analyzed by mass spectrometry to quantify drug levels in both plasma and tumor tissues.

### Immunohistochemistry

For the IHC studies, GBM tissue specimens from the brains of euthanized mice were fixed in formalin and embedded in paraffin-embedded (FFPE) tissue blocks. Then, the serial unstained slides were cut from these FFPE tissue blocks to stain for the infiltrating immune cell marker after treatment with f-MET and P4-MET. IHC for CD8a (#Cat-ab209775), CD4 (#Cat-ab183685), Treg (#Cat-ab36607), CD68 (#Cat-ab125212), and CD206 (#Cat-ab64693) was performed using the Benchmark IHC/ISH system (Roche, Basel, Switzerland) after incubating the slides with each antibody separately. All these antibodies were purchased from Abcam (Cambridge, MA, USA). After staining with these markers, the slides were scanned at 40x magnification using the Ventana iScan HT slide scanner. Quantification was performed using Definiens Tissue Studio (Ventana, Munich, Germany).

### Histological analysis

Histological analysis was performed to assess potential tissue damage or toxicity induced by f-MET and P4-MET following *in vivo* administration. Histopathological evaluation was conducted on key organs, including the brain, to examine any signs of organ-specific toxicity, inflammatory response, or structural abnormalities that could result from systemic administration of the formulations. Mice were euthanized at predetermined time points after receiving f-MET or P4-MET treatments, and tissue samples were harvested for subsequent histological examination. The tissues were fixed in formalin, embedded in paraffin, and sectioned for hematoxylin and eosin (H&E) staining to evaluate general tissue morphology. The histological sections were then analyzed for signs of toxicity, including inflammation, necrosis, fibrosis, and cell infiltration, particularly in the liver and kidneys, the primary organs for detoxification. The brain tissue was examined for evidence of tumor growth with assistance from a neuropathologist.

GBM tumor-bearing mouse CT-2Aluc cells were intraperitoneally injected with P4-MET-IR800 (200mg/kg based on free metformin) once. After 8 hours, the mouse was sacrificed, and the brain, liver, and pancreas were quickly embedded in OCT (optimal cutting temperature) and cooled by dry ice. The tissue sections were then cut to a thickness of 6-10 μm using the cryostat at -20 °C, fixed, and stained with DAPI. Cryosections were visualized by a confocal laser scanning microscope (Zeiss Cell Discoverer 7).

### Flow cytometry

Single-cell suspensions were prepared from collected tissue samples and stained with fluorochrome-conjugated monoclonal antibodies targeting surface and intracellular T cell markers. The following antibodies were used: CCR7-BUV661, CD3-EF506, CD4-BUV395, CD8-Super Bright 600 (SB600), CD11b-PE-Cy5, CD25-BV711, CD45-PE-Cy7, CD45RA-BV421, CD45RO-BV750, and FoxP3 Alexa Fluor 488. Dead cells were excluded using Fixable Viability Dye UV Blue. All antibodies were used according to the manufacturer’s instructions. Surface staining was performed for 30 minutes at 4°C in FACS buffer (PBS supplemented with 2% FBS and 2 mM EDTA). For intracellular staining of FoxP3, cells were fixed and permeabilized using the FoxP3/Transcription Factor Staining Buffer Set, then stained in the permeabilization buffer for 30-45 minutes at room temperature.

Data acquisition was performed using a BD LSRFortessa X-50 flow cytometer (BD Biosciences), equipped with five lasers (355 nm, 405 nm, 488 nm, 561 nm, and 640 nm) and capable of detecting up to 50 parameters. Single-stained compensation controls and fluorescence-minus-one (FMO) controls were included for accurate gating and compensation. At least 100,000 live events were collected per sample. Data was analyzed using FlowJo software (v10.x, BD Biosciences).

### RNA Sequencing (RNA-Seq) Analysis

Total RNA was isolated from U87 glioblastoma cells following treatment with 1 mM f-MET and an equivalent dose of P4-MET at 4 h, 24 h, and 48 h using the RNeasy Mini Kit (Qiagen, Hilden, Germany) according to the manufacturer’s instructions. RNA concentration and purity were assessed using a NanoDrop spectrophotometer (Thermo Fisher Scientific, Waltham, MA, USA). Extracted RNA samples were stored at −80 °C until further processing. RNA integrity and quality were evaluated using the Bioanalyzer 2100 system (Agilent Technologies, Waldbronn, Germany). For RNA sequencing, 200 ng of total RNA per sample was used for library preparation and downstream transcriptomic analysis.

### Differential Gene Expression Analysis

Differential gene expression analysis was performed using the DESeq2 package in R to identify transcriptomic alterations induced by f-Met and P4-MET relative to untreated control samples. Gene expression changes were quantified as log2 fold changes (Log2FC) relative to control conditions at three independent time points (4 h, 24 h, and 48 h). Statistical significance was determined using adjusted p-values (padj) generated by the Benjamini–Hochberg false discovery rate (FDR) correction method.

### Gene Selection and Filtering Criteria

A comparative analysis was performed to identify genes that show strong responsiveness to f-MET but reduced or attenuated responsiveness to P4-MET treatment. The candidate genes were filtered based on the following criteria: high induction in the f-MET condition (Log2FC > 1.0), statistical significance (adjusted p-value, padj < 0.05), and low induction in the P4-MET-treated condition (Log2FC < 1.0). The top 10 genes from each time point were selected for downstream comparative visualization and interpretation.

### Statistical Visualization and Bioinformatic Analysis

Data visualization and statistical analyses were performed in R using the ggplot2, dplyr, and tidyr libraries. Comparative bar plots were generated to visualize Log2FC values for f-Met and PMET treatments side by side for each selected gene.

### Software and Script Implementation

All transcriptomic analyses and visualizations were implemented using custom R scripts, including finding upregulated genes in f-MET vs downregulated genes in P4-MET. All analyses have been used by the DESeq2 package for calculation and normalization.

### siRNA Transfection and Bola2b Knockdown

Primary human GBM Cells (GBM22) were transfected with either 10 nM or 25 nM Bola2b siRNA SMARTpool (Dharmacon, Lafayette, CO, USA) or 25 nM control siRNA-A (sc-37007; Santa Cruz Biotechnology, Dallas, TX, USA). In parallel, pharmacological inhibition studies were conducted using 50 μM and 100 μM deferoxamine (DFO; Santa Cruz Biotechnology), a functional inhibitor associated with BOLA2B-related iron metabolism pathways.

Transfections were performed using TurboFect Transfection Reagent (Thermo Fisher Scientific) according to the manufacturer’s instructions. GBM22 cells were seeded in 6-well plates and transfected in Opti-MEM reduced-serum medium. After 6 h of incubation, the transfection medium was replaced with complete DMEM containing fetal bovine serum and antibiotics. At 48 h and 72 h post-transfection, total RNA and protein lysates were collected, respectively, for downstream validation analyses. Reverse transcription quantitative PCR (RT-qPCR) and immunoblotting assays were performed to confirm the efficiency of BOLA2B knockdown at both the transcript and protein levels.

### Statistical analysis

Results are presented as mean ± SD. A two-sided Student’s T-test was conducted to assess the statistical significance of the differences between the two groups. Two-way ANOVA was used to determine differences among multiple groups. P < 0.05 was considered statistically significant. The K-M curve was analyzed for overall survival in each treatment group.

## Results

### Synthesis and characterization of P4-MET

We successfully synthesized and preliminarily characterized P4-MET, a metformin-conjugated dendrimer formulation in which metformin is covalently attached to P4-OH [Generation 4, G4(OH)₆₄], a polyamidoamine dendrimer with an ethylenediamine core and terminal hydroxy amidoethanol surface groups. The P4-MET conjugate was synthesized via a two-step reaction: first, activation of dendrimer surface hydroxyl groups using 1,1’-carbonyldiimidazole (CDI) to form reactive intermediates; second, coupling with free-base metformin to form carbamate linkages (**Fig. 1A**). The resulting P4-MET conjugates were purified by dialysis and Sephadex LH-20 size-exclusion chromatography, followed by analysis using HPLC to ensure removal of unreacted metformin and reaction byproducts.

**Figure 1.**
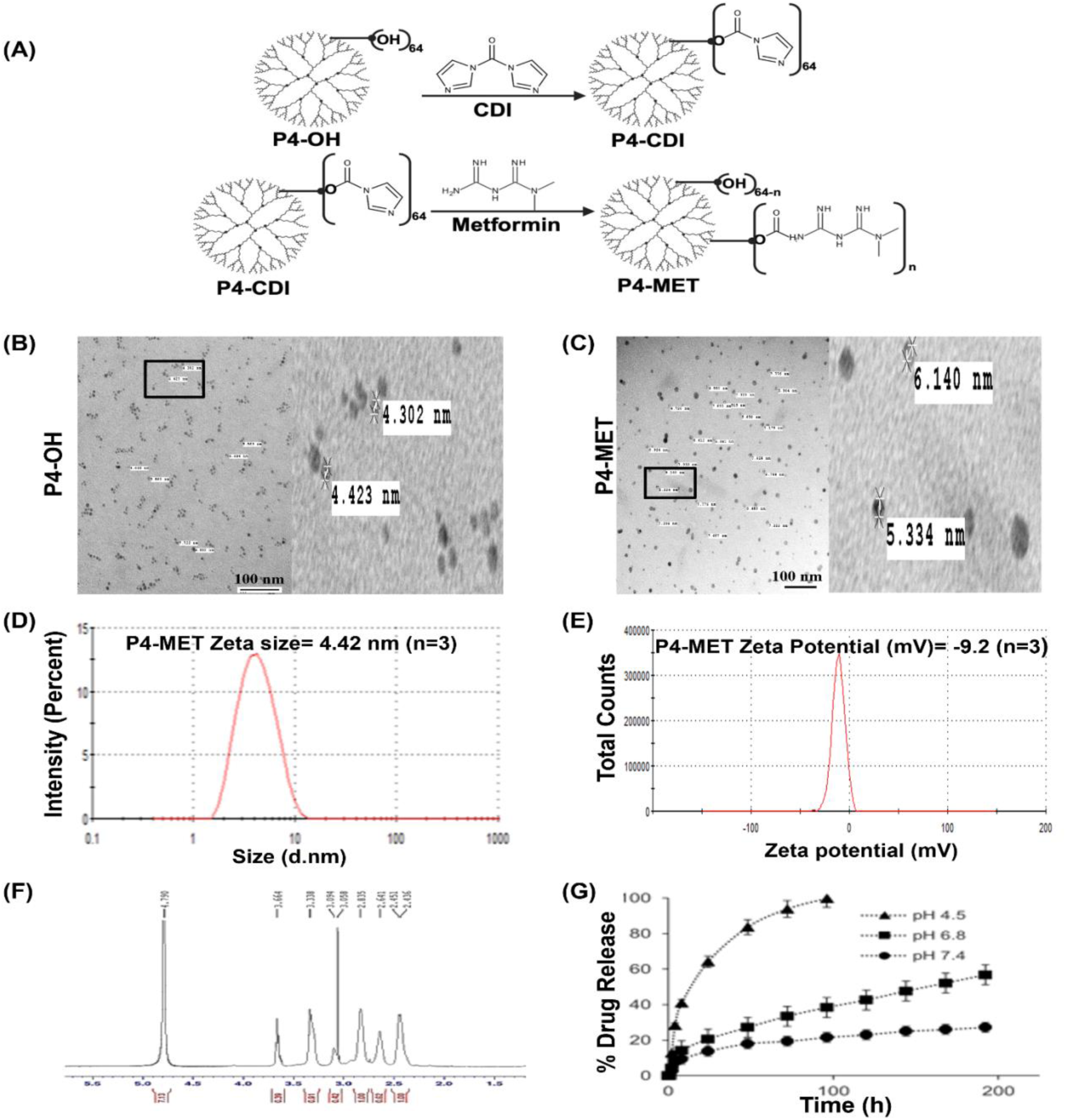
Synthesis and physicochemical characterization of P4-MET. (**A**) Schematic illustration of P4-MET synthesis. Free base metformin was conjugated to the hydroxyl group of a PAMAM dendrimer via a cleavable carbamate bond. (**B, C**) TEM characterization of P4-OH and P4-MET. Samples were prepared in methanol at a concentration of 5 mg/mL, dropped-cast onto carbon-coated copper grids, and air-dried before imaging. (**D**) The hydrodynamic size and (**E**) the zeta potential of P4-OH (vehicle) and P4-MET were measured by dynamic light scattering (DLS) using a Zetasizer Nano ZS (Malvern Instruments). Samples were prepared in methanol at a concentration of 5 mg/mL and analyzed at 25 °C. Measurements represent the average of three independent runs. (**F**) ¹H NMR spectrum of the P4-MET was recorded in D_2_O at 400 MHz. (**G**) In vitro release profile of free metformin from P4-MET at different pH, simulating relevant biological conditions: 4.5 (lysosomal), 6.8 (tumor microenvironment), and 7.4 (physiological).

The ¹H-NMR spectrum confirmed the successful conjugation of metformin to the P4-OH dendrimer. Key proton signals from metformin, such as the N–CH₃ singlet (∼δ 2.9 ppm) and other methylene and amidine protons (∼δ 3.0–5.0 ppm), were observed in the conjugate spectrum, with slight chemical shift changes relative to free metformin, indicating covalent modification rather than simple physical adsorption. The characteristic broad peaks of the PAMAM backbone, including methylene protons of the dendritic core and branches (∼δ 2.2–3.5 ppm), were retained (**Fig. 1F**). Integration of metformin signals relative to PAMAM methylene peaks enabled estimation of the average degree of substitution (∼20–25 metformin molecules per dendrimer). Based on batch-to-batch analysis, the drug loading capacity was consistently around 30% w/w, corresponding to approximately 20–25 metformin molecules conjugated per dendrimer (**Fig. 1F**). DLS analysis demonstrated that the P4-MET particles have a narrow size distribution, with a median hydrodynamic diameter of ∼5 nm, consistent with the PAMAM G4 core plus surface modification. The polydispersity index (PDI) was low (< 0.2), indicating a uniform nanoparticle population. Small particle size and near-neutral zeta potential (-9.2 mV), characteristics favorable for cellular uptake and membrane penetration, similar to the unmodified PAMAM-OH dendrimer backbone (**Fig. 1D and E**). TEM imaging revealed that P4-MET nanoparticles are uniformly spherical, with a well-dispersed morphology and no visible aggregation (**Fig. 1C**). The introduction of carbamate bonds enables acid-sensitive drug release, promoting preferential metformin release under acidic conditions, such as those found in intracellular endosomes or tumor microenvironments (**Fig. 1G**).

Furthermore, the endotoxin limit (EL) calculation was done by two different methods and the results showed that the endotoxin limit of P4-MET (NCL517-1) was 0.12 EU/mg which is below the EL (This endotoxin limit is applicable at an HED of 40.65 mg/kg as per NCL report) and no microbial contamination was detected in the sample (**Table 1, and Table 2**).

**Table 1:** Endotoxin Limit Calculation.

| **P4-MET (NCL517-1)** |  |
| --- | --- |
| Sample Concentration (mg/ml of metformin) | 500 |
| Sponsor's Mouse Dose (mg/kg) | 500 |
| Human Equivalent Dose [HED] (mg/kg) | 40.65 |
| Theoretical Human Plasma Concentration at HED [THPC] (mg/ml) | 0.5 |
| **Endotoxin Limit [EL] (EU/mg)** | **0.12** |
| **Maximum Valid Dilution (MVD) for LAL Assays** |  |
| MVD for Turbidity LAL | 61500-fold |
| MVD for Chromogenic LAL | 61500-fold |
| MVD for Gel-Clot LAL | 2050-fold |

**Table 2:**
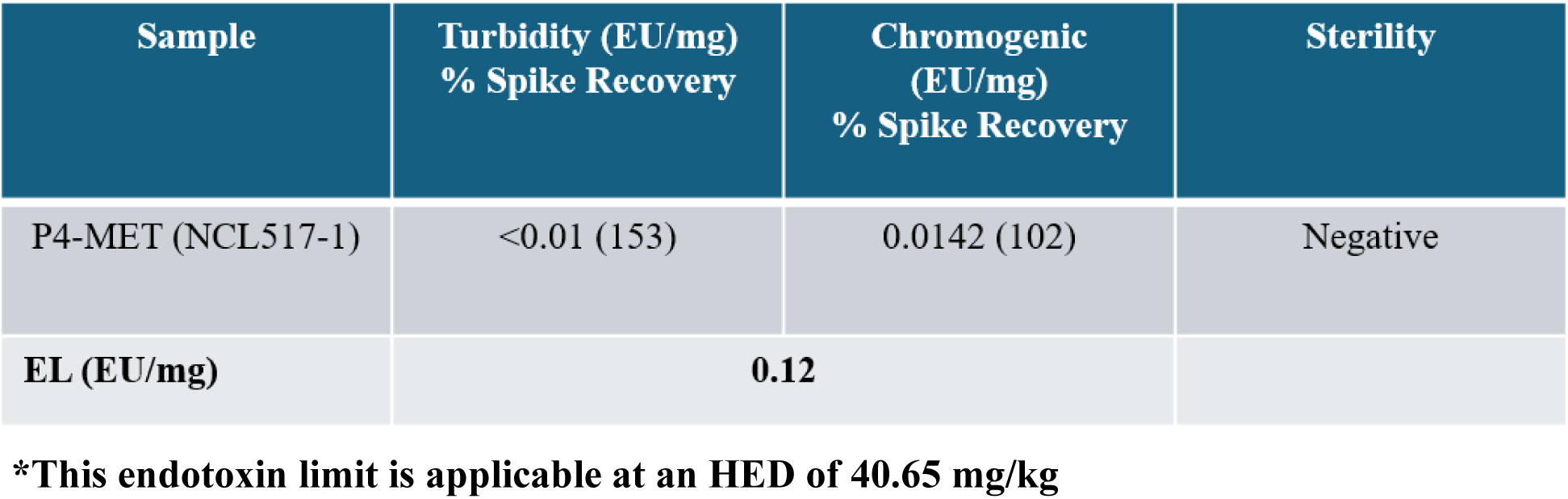
Endotoxin Limit.

### Enhanced anti-proliferative effects of P4-MET over free metformin (f-MET) in adult and pediatric glioblastoma cells

We evaluated the effects of P4-MET and f-MET on a panel of glioblastoma cell lines using the CTB assay. This panel included adult human GBM lines U87MG (**Fig. 2A**) and LN229 (**Fig. 2B**), mouse glioma lines CT-2Aluc (**Fig. 2C**) and MGPP3 (**Fig. 2D**), as well as the pediatric high-grade glioma (HGG) cell lines DMG4423 (**Fig.2E**) and SF188-luc (**Fig.2F**), and a patient-derived primary GBM cell line (**Fig. 2G**), established under an IRB-approved protocol at our institution. All tested GBM cell lines exhibited dramatically enhanced inhibitory effects on cell proliferation with P4-MET treatment, with 50% inhibitory concentrations (IC₅₀) improved 50-250-fold over f-MET, except for LN229, which was relatively insensitive to f-MET. Treatment with P4-OH alone, used as a control, showed no toxicity in any of the cell lines tested. Moreover, the simple combination of P4-OH and free metformin (f-MET + P4-OH) did not enhance efficacy compared to free metformin alone, underscoring the importance of covalent conjugation for improved cell penetration and drug efficacy (**Fig. 2**). Beside this we also performed the trypan blue exclusion assay to evaluate the comparative cell death between GBM cells (U138), and normal human cortical neurons (HCN-2) after the P4-MET treatment and found significant dose-dependent cell death in U138 but not in HCN-2 at the same concentration, suggesting a tumor-specific cytotoxicity (Data not shown).

**Figure 2.**
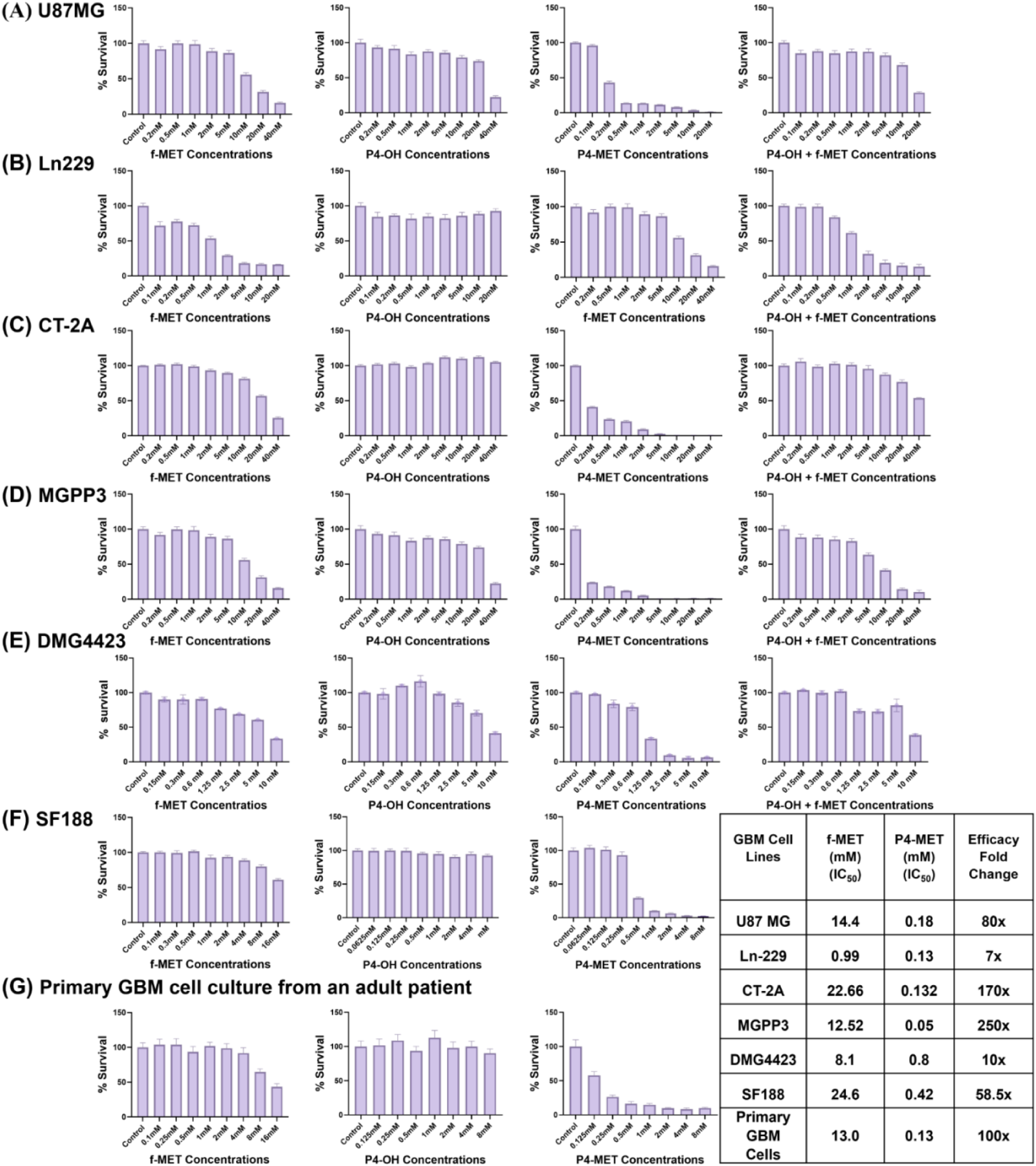
Significantly enhanced inhibitory effects on cell proliferation by P4-MET compared with f-MET on (**A**) U87MG (adult human GBM), (**B**) Ln229 (adult human GBM), (**C**) CT-2Aluc (mouse GBM), (**D**) MGPP3 (mouse GBM), (**E**)DMG4423 (pediatric GBM), **(F)** SF188 (human pediatric GBM), and (**G**) primary GBM cells isolated from an adult GBM patient cultured *in vitro*. Cells were treated with free metformin (f-MET), P4-OH, P4-OH mixed with free metformin (P4-OH + f-MET), and P4-MET for 72 hours with indicated concentrations equivalent to free metformin content. Cell proliferation was estimated using the Cell Titer Blue assay. The IC_50_ for the GBM cell line (U138MG), tested in the lab but not shown here, exhibited similar trends. All experiments were performed with six independent replicates. The P4-OH + f-MET combination treatment was not performed for (**F**) SF188^luc^ and (**G**) primary GBM cells.

### Intracellular uptake of P4-MET demonstrated by confocal microscopy and transmission electron microscopy (TEM)

The cellular uptake of P4-MET was examined by confocal microscopy at different time intervals (4 hours, 16 hours, and 24 hours) after treating CT-2Aluc cells with A488-tagged P4-MET. The results showed time-dependent accumulation of the drug in the cells. The fluorescence from the tagged drug begins to show as early as 4 hours after treatment, indicating significant accumulation in GBM cells (**Fig. 3A**). Additionally, the results demonstrate substantial in vitro cellular uptake of A488-tagged P4-MET in bone marrow-derived macrophages (BMDM) after 24 hours of treatment, targeting mitochondria (**Fig. S1**).

**Figure 3.**
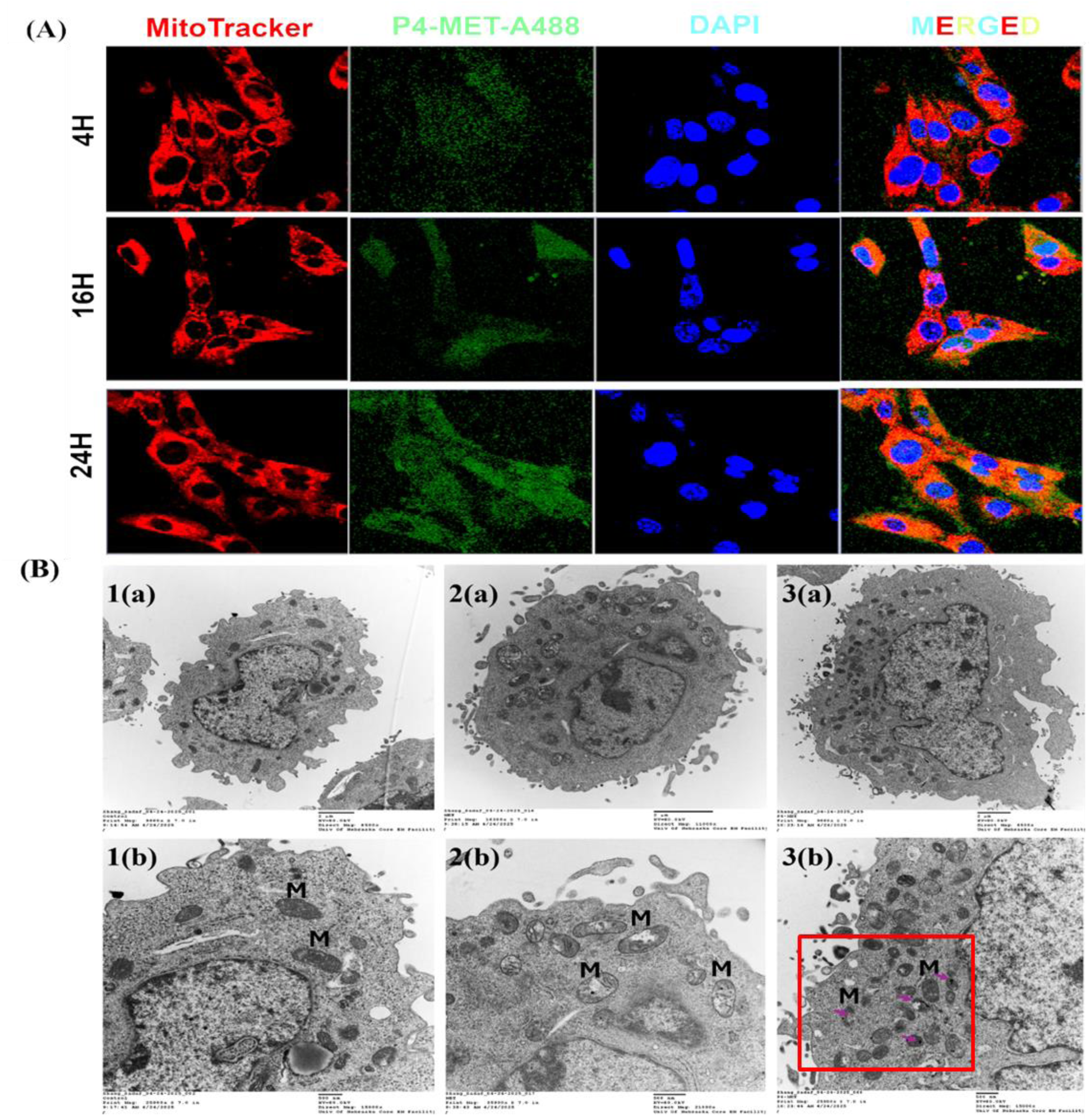
Confocal microscopy demonstrates intracellular uptake of P4-MET by CT-2Aluc cells. Detectable intracellular P4-MET tagged with A488 can be detected as early as 4 hours after incubation (**A**). Transmission electron microscopy of CT-2Aluc cells showed mitochondrial structural changes (**B**): (**1**) Control cells without treatment; (**2**) f-MET (1mM) treated CT-2Aluc cells; and (**3**) P4-MET (1 mM, equivalent to f-MET) treated CT-2Aluc cells. Arrows indicate the alterations in mitochondrial morphology and drug bioaccumulation. (Panel ‘a’ corresponds to 2µm, and panel ‘b’ corresponds to a 500nm scale bar in each group). M: mitochondria.

To further evaluate the ultrastructural changes and drug bioaccumulation, we analyzed CT-2Aluc cells by TEM after 24 hours of treatment with 1 mM f-MET and an equivalent dose of P4-MET. The P4-MET treated cells showed significantly altered mitochondrial structure and accumulation of the drug in the mitochondria (**Fig. 3B, 3a, and b**) compared to untreated CT-2Aluc cells, which showed normal intact mitochondrial structure (**Fig. 3B, 1a, and b**). After the f-MET treatment, most of the mitochondria were swollen up (**Fig. 3B, 2a, and b**), though the effect was more pronounced in the case of P4-MET treatment.

### Immunoblotting assays demonstrate differential regulation in AKT-mTORC signaling pathways between f-MET and P4-MET

Immunoblotting was performed to evaluate the effects of P4-MET versus f-MET on key signaling pathways involved in cell growth and proliferation in CT-2Aluc mouse glioblastoma cells. Both f-MET and P4-MET treatments showed time-dependent upregulation of AMPK levels (**Fig. 4**) due to energetic stress resulting from reduced oxidative phosphorylation-mediated mitochondrial ATP production and a concurrent increase in glycolytic ATP production (**Fig. S2**). Interestingly, we observed opposite responses in several molecular markers when comparing the two treatment groups. Specifically, P4-MET treatment resulted in a marked reduction in the phosphorylation of ribosomal protein S6 at Ser235/236 (p-S6 235/236), a downstream target of the mTORC1 signaling pathway, while total S6 protein levels were modestly decreased. In contrast, f-MET treatment did not decrease p-S6 levels and, in some cases, appeared to slightly increase them, suggesting a limited impact on mTOR activity in this context.

**Figure 4.**
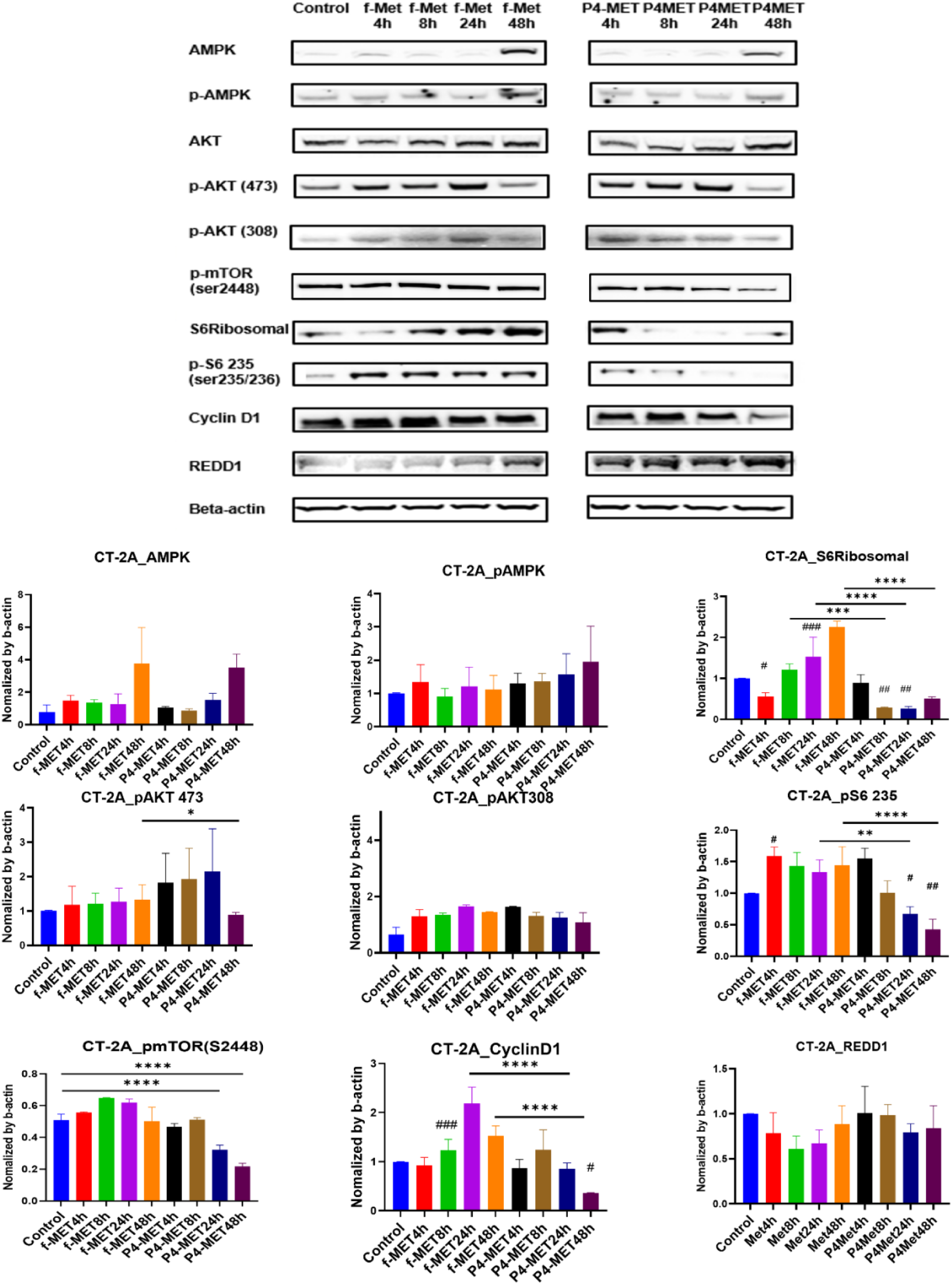
Immunoblotting shows the effects of 1 mM f-MET and an equivalent dose of P4-MET on different signaling pathways in CT-2Aluc cells after 4h, 8h, 24h, and 48h of treatment.

Similarly, Cyclin D1, a key regulator of cell cycle progression, was significantly downregulated in response to P4-MET, but remained largely unchanged following free metformin treatment. Notably, mTOR phosphorylation at Ser2448, a hallmark of mTORC1 activation, was also reduced upon P4-MET exposure but not affected by f-MET, further supporting the notion that P4-MET more effectively inhibits mTORC1 signaling (**Fig. 4**).

### Identification of Bola2b as the top differentially regulated gene between f-MET and P4-MET treatment in GBM cells

Following RNA-Seq and differential expression analyses comparing f-MET- and P4-MET-treated U87MG cells, BOLA2B was identified as one of the most significantly differentially expressed genes 4 hours after drug treatment. f-MET treatment induces significant upregulation in BOLA2B mRNA expression compared to P4-MET (**Fig. S5A**).

To further investigate the molecular mechanisms associated with drug efficacy and the functional role of BOLA2b, a knockdown experiment was performed in the GBM22 patient-derived xenograft (PDX) glioblastoma cell line.

The correlations between mTORC1 and BOLA2B, and between mTORC1 and GPX4 (a Ferroptosis inhibitor) were investigated in TCGA. Interestingly, we found BOLA2B is positively correlated with mTORC1 and GPX4 (**Fig. S5D**). These findings prompted us to consider the potential role of ferroptosis, a cell death mechanism, in GBM cells following P4-MET treatment. As BOLA2B is a critical modulator of iron metabolism and mTORC1 activity in GBM, we further evaluate the p-mTORC1 level after BOLA2B knockdown in GBM cells. The qRT-PCR data confirmed a decrease in BOLA2B expression in GBM 22 cells in both the siRNA-transfected and DFO-treated groups compared with the control group (**Fig. S5E**). Additionally, immunoblotting results also showed downregulation of p-mTORC1 expression compared to control following significant knockdown of BOLA2B at 25nM, further confirming the positive correlation (**Fig. S5F**). Additionally, significant downregulation in the p-mTORC1 level in CT-2A GBM cells (**Fig. 4**) and an increase in the number of small, condensed mitochondria with reduced cristae features, as shown in TEM images (**Fig. 3B**) after the P4-MET treatment, as compared to f-MET, indicate the P4-MET treatment might induce the susceptibility of GBM cells to ferroptosis-mediated cell death, which may be protected by f-MET treatment by significantly promoting BOLA2B expression.

These findings suggest that P4-MET achieves enhanced intracellular delivery and mechanistic engagement compared with f-MET, potentially due to improved cellular uptake and sustained intracellular retention. The differential regulation of mTOR pathway components further highlights the advantage of the dendrimer-based delivery system in modulating cancer-relevant signaling pathways more effectively than the unconjugated drug.

### P4-MET, but not f-MET, significantly improved the overall survival (OS) in an orthotopic syngeneic adult GBM mouse model when used alone or combined with brain RT, with no detectable acute toxicities

Orthotopic syngeneic mouse GBM models have been established using the stereotactic microinjection technique routinely performed in our laboratory (**Fig. 5A**). We implanted 5 x 10^4 CT-2Aluc cells, a GBM cell line stably transfected with a luciferase reporter gene, into the right hemisphere. P4-MET and f-MET (200mg/kg/d equivalent to free metformin) daily IP injection started on day 8 post-implantation for 3 weeks with or without whole brain RT with a total dose of 8Gy in one fraction on day 11 (**Fig. 5B**). Overall survival (OS) of each group of mice (8-12 mice per group) was compared. P4-MET treatment alone, but not f-MET alone, prolonged OS of the tumor-carrying mice (orange line in **Fig. 5C**), and so did RT alone (red line). Free metformin sensitized RT with moderate OS improvement compared with either treatment alone (purple line). Impressively, we observed a significantly stronger synergistic effect when combining P4-MET and RT (black line in **Fig. 5C**) than when combining f-MET and RT. After tagging P4-MET with the infrared fluorescent dye IR800 (Li-Cor) for injection and live whole-body fluorescent scanning using the Pearl Trilogy Imaging System (Li-Cor, Lincoln, NE), we also studied the pharmacodynamics of P4-MET penetration into tumor-bearing tissue (brain). Significantly increased tumor penetration and prolonged tissue retention of the drug were observed when RT was delivered simultaneously with P4-MET injection with Tissue/background ratio (TBR) of detectable fluorescent signals in tumor-bearing brain reached higher peak (purple line) than no RT (green line) or no tumor/normal brain (red line) controls (**Fig. 5D**). Without RT, fluorescent signal decreased soon back to background within one day but persisted at 3-4 x of background for days post-RT/drug injection.

**Figure 5.**
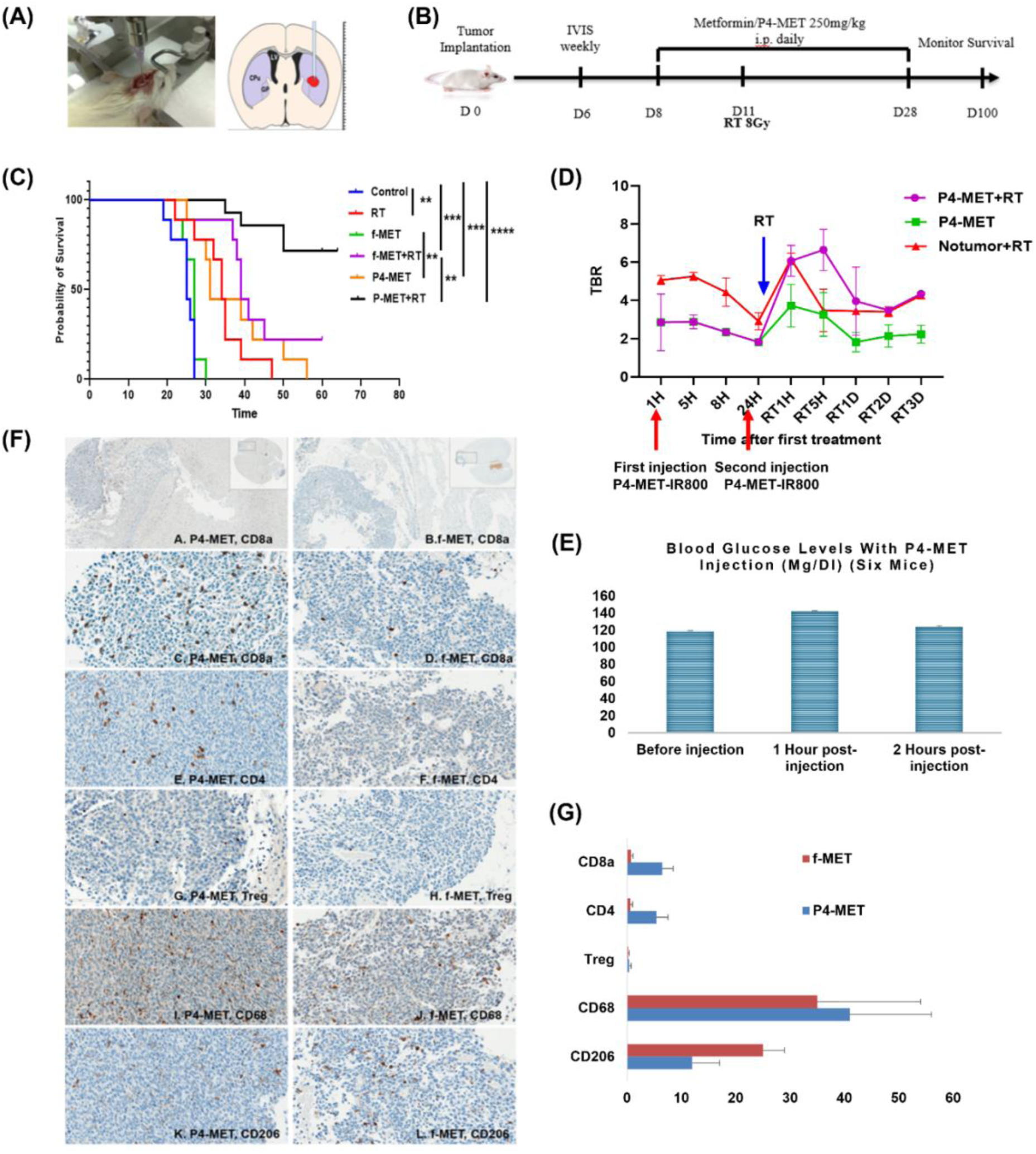
*In vivo* studies on mice with orthotopic GBM tumor implant after treatment. (**A**) Stereotactic microinjection of CT-2Aluc mouse GBM cells to the right hemisphere of mice brain. (**B**) Treatment scheme after confirming brain tumor growth with weekly IVIS. (**C**) Kaplan-Meier’s OS curves of mice treated with free metformin (f-MET), P4-MET, with or without brain RT. (**D**) Fluorescence-tagged P4-MET (P4-MET-IR800) showed increased brain penetration when combined with RT. (**E**) Stable blood glucose levels before and after P4-MET injection. (**F**) IHC of mouse brain tissues stained for CD8a, CD4, Treg, CD68 T cell markers and CD206 (M2Ø) after P4-MET treatment or free met. Marker stained: Brown; Nuclei: blue. (**G**) Quantification of infiltrating tumor cells density in the tumor microenvironment (TME).

Furthermore, Immunohistochemistry (IHC) studies demonstrated significantly increased CD8+ cytotoxic T cell infiltration in P4-MET-treated brain tumors in mice, with positive staining density approximately 8× higher than in f-MET-treated tumors (**Fig. 5F and G**). Staining for CD206, an M2 macrophage marker, in the same tumors showed a significant decrease in staining intensity with P4-MET treatment, indicating greater efficacy of P4-MET in suppressing the immunosuppressive TAM subtype *in vivo*. No significant acute/subacute toxicities were identified in mice treated with P4-MET, including weight, comprehensive biochemical panel (CMP), and complete blood counts (CBC) in peripheral blood tests (data not shown), as in pre- and post-injection blood glucose levels (**Fig. 5E**) in P4-MET treated and tumor-controlled mice compared with healthy mice. No hypoglycemia was observed during the first two hours after P4-MET injection in any of the six mice tested (**Fig. 5E**). Histological studies were also performed on the dissected normal organ tissues including heart, liver, lung, pancreas, spleen, small intestines, and muscle in four mice treated with P4-MET with no abnormalities including necrosis, thrombosis, etc., were detected (data not shown). Only one mouse treated with f-Met developed conjunctivitis, which was successfully treated with topical antibiotics.

### P4-MET preferentially accumulates in tumor-bearing brain tissue, particularly after irradiation

The P4-MET conjugate preferentially accumulates in tumor-bearing brain tissue due to the unique characteristics of the P4-OH dendrimer and its ability to penetrate the BBB upon systemic administration. The P4-MET formulation leverages enhanced permeability and retention (EPR) effects, where the leaky vasculature characteristic of tumors facilitates the accumulation of nanoparticles within the tumor microenvironment. This increases local drug concentration at the tumor site, ensuring more effective targeting of the tumor cells. Moreover, the small size and hydrophilic nature of P4-OH facilitate crossing the blood-brain barrier, thereby enhancing the direct delivery of metformin to the tumor. In vivo imaging studies (**Fig. 6A**) and histological analysis (**Fig. 6C, a-d**) confirmed the preferential accumulation of P4-MET in GBM tumors, with minimal off-target distribution to healthy brain tissue or other organs. This selective accumulation in the tumor microenvironment suggests that P4-MET holds significant promise as a targeted therapeutic strategy for glioblastoma treatment, potentially improving drug efficacy while reducing systemic side effects (**Fig. 6**). Further, LC/MS quantification showed significant accumulation of f-MET in the mouse plasma (**Fig. 6B, a-d**) and liver (**Fig. 6B, e-h**) in a time-dependent manner after the last injection of P4-MET. The mouse brain with a tumor showed a remarkable accumulation of f-MET after P4-MET combined with RT treatment at 6 hours and 24 hours post-treatment (**Fig. 6D**).

**Figure 6.**
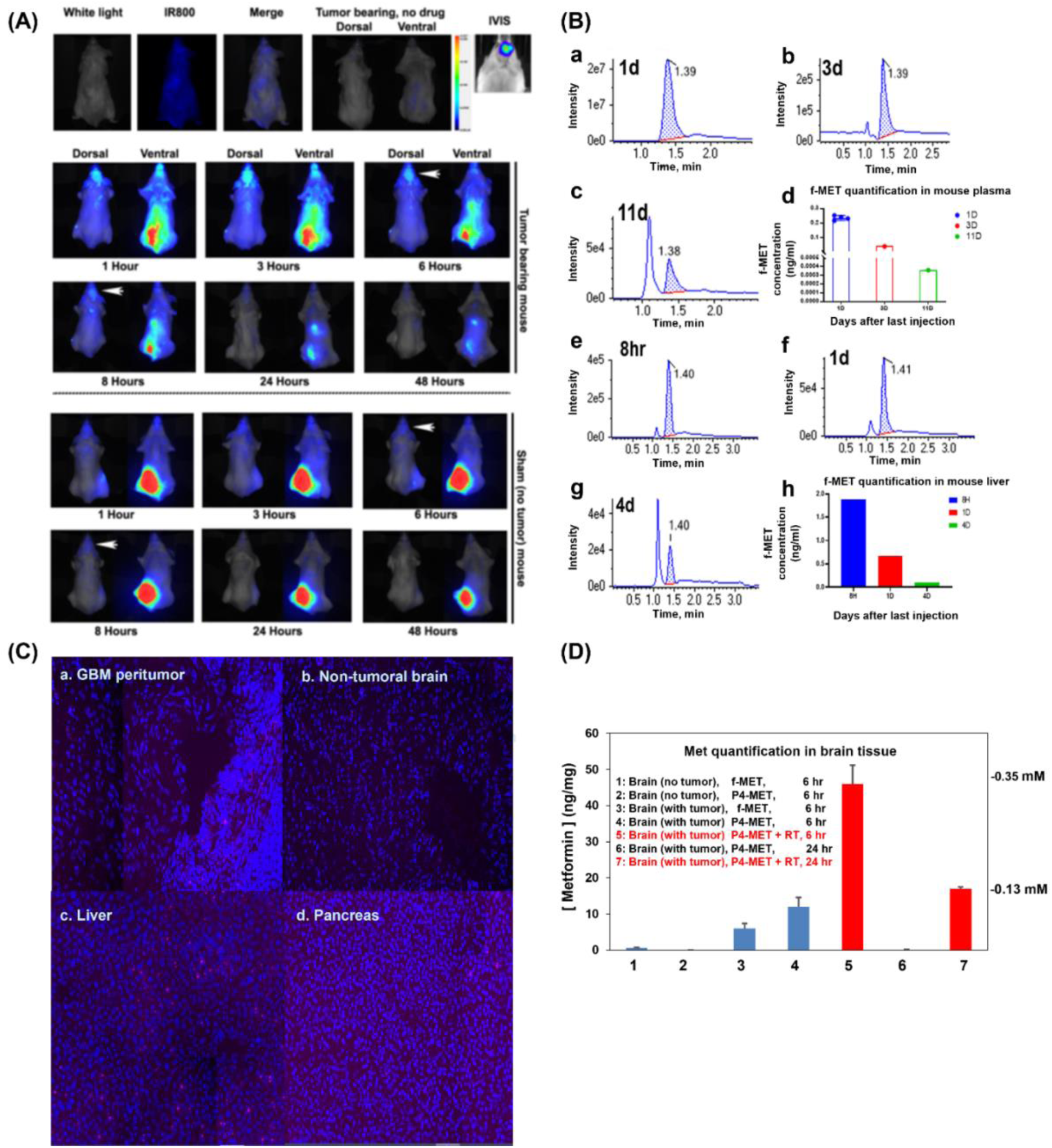
Tumors in the brain significantly increase drug accumulation compared with the non-tumor-bearing mouse brain (sham with cranial surgery but no tumor implant) (white arrows). (**A**) Fluorescent-labelled drug became barely detectable in brain after 24-48 hours. P4-MET showed significantly higher concentrations in tumor-bearing brain than free metformin (f-MET) and RT to brain significantly prolonged the drug accumulation in the tumor-bearing brain. (**B**) Quantification of f-MET in mouse plasma and liver tissue by LC/MS. Metformin-d6 (Item #16921, Cayman Chemical) was used as an internal standard. Albino B6 female mice were treated with f-MET (200mg/kg, daily i.p. injection) for two weeks. Plasma and liver tissues were collected to quantify drug concentrations at the indicated hours/days after the last dose of injection. (**B, a-d**): LC/MS analyses of f-MET in mouse plasma specimens that were collected 1 day (**a**), 3 days (**b**), and 11 days (**c**) after the last IP injection of f-MET. (**d**) Comparison of the quantified metformin concentrations in mouse plasma based on the calibration curve (not shown). (**B, e-h**) LC/MS analyses of f-MET in mouse liver tissues that were collected 8 hours (**e**), 1 day (**f**) and 4 days (**g**) after the last IP injection of f-MET. (**h**) Comparison of the quantified metformin concentrations in mouse liver tissues. (**C**) Histological analysis (**a-d**) confirmed the preferential accumulation of P4-MET in GBM tumors (**a**), with minimal off-target distribution to healthy brain tissue or other organs. (**D**) LC/MS analysis of metformin concentrations in the brain (no tumor) or brain tumor tissues after f-MET and P4-MET IP injection with and without RT.

### P4-MET promotes effector memory T cells (T_EM) in systemic immunity

The P4-MET conjugate modulates the immune response in GBM tumor-bearing mice by promoting the expansion of effector memory T cells (T_EM cells), which play a crucial role in adaptive immunity. Effector memory T cells are characterized by their ability to rapidly respond to previously encountered antigens, making them essential for sustained immune surveillance and long-term tumor control. Only one out of the four mice that survived 4 months after initial P4-MET and brain RT treatment developed tumors after rechallenging with CT-2Aluc cells implanted the second time (**Fig. 7B**). All eight treatment naive mice developed brain tumors (per IVIS imaging) after CT-2Aluc GBM cells implanted as controls (**Fig. 7A**). Results indicate formation of long-term memory of immunity after P4-MET+RT treatment.

**Figure 7.**
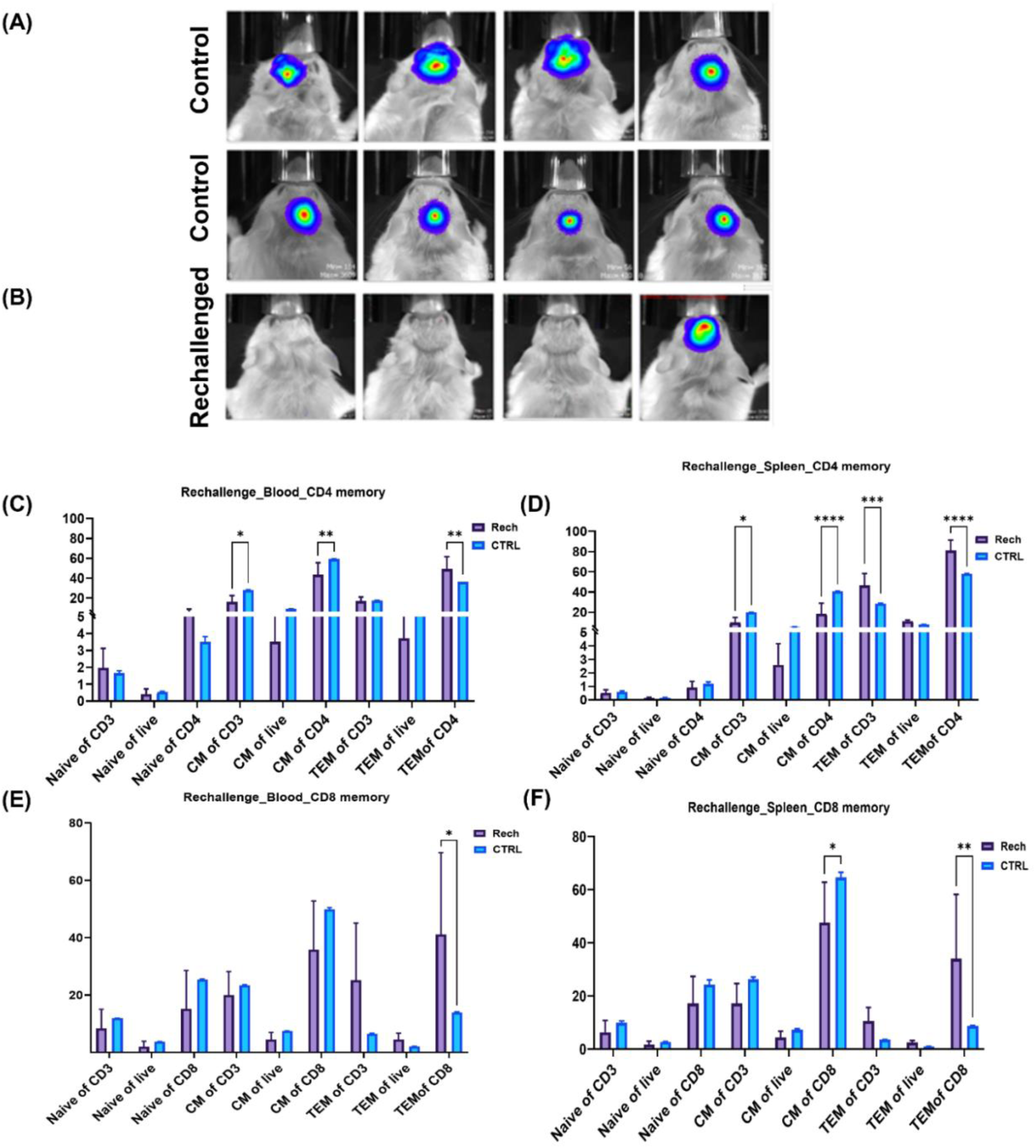
MET plus RT treatment leads to long-term immune memory. (**A**) All eight control mice developed a brain tumor (per IVIS imaging) after CT2A GBM cells. (**B**) Only one out of the four rechallenged mice developed tumors after initial P4-MET and brain RT treatment. Upon treatment with P4-MET, there was a significant increase in the frequency and activation of T_EM cells, as indicated by CD4 (**C and D**) and CD8 (**D and E**) analysis in the peripheral blood and spleen.

Upon treatment with P4-MET, there was a significant increase in the frequency and activation of T_EM cells of CD4 (**Fig. 7C and D**) and CD8 (**Fig. 7E and F**) within the peripheral blood and the spleen. This expansion of T_EM cells was associated with increased immune infiltration into the tumor, suggesting that P4-MET not only targets the tumor directly via its drug-delivery system but also modulates the immune landscape in favor of anti-tumor immunity.

Besides this, other memory immune cells also showed enhanced activation followed by P4-MET+RT treatment as compared to other treatment groups in the blood samples (**Figure S3, C and D**) and in spleen samples (**Figure S4, C and D**), depicting P4-MET mediated radio-sensitization in GBM.

## Discussion

The synthesis of P4-MET yielded a nanocarrier approximately 5 ± 2 nm in size, consistent with P4-OH dendrimers, which typically range from 4 to 5 nm in diameter. The small size of P4-MET is a critical attribute, as it enables greater cellular uptake and efficient tissue penetration, both of which are essential for the effective delivery of therapeutic agents to target cells. The nanoscale size enhances P4-MET’s potential to accumulate in tumor tissues via the enhanced permeability and retention (EPR) effect, a phenomenon commonly observed in tumor environments due to leaky vasculature. The release profile exhibited an initial burst phase followed by a sustained release phase. The burst release was more pronounced at pH 5.5, indicating pH-sensitive release behavior. This enhanced release in acidic conditions suggests that P4-MET effectively exploits the tumor microenvironment to facilitate drug release at the target site, thereby improving therapeutic efficacy while minimizing off-target effects. These findings highlight the potential of P4-MET as a tumor-targeting drug delivery system with enhanced brain penetration and controlled drug-release properties, making it a promising candidate for the treatment of glioblastoma^18, 19^.

The zeta potential of measured P4-MET was -9.2 ± 2 mV, suggesting a relatively stable formulation. The positive charge of the dendrimer surface provides electrostatic repulsion between particles, minimizing aggregation and ensuring colloidal stability in solutions. A low polydispersity index (PDI) of 0.23 indicates that the particles exhibit a narrow size distribution, a desirable feature for consistent drug delivery performance. Furthermore, the metformin loading capacity of P4-MET was approximately 20%, indicating efficient drug encapsulation and a promising drug-to-carrier ratio suitable for achieving therapeutic effects at relatively low doses. These results underscore the successful development of a stable, homogeneous, and drug-loaded nanocarrier with favorable characteristics for drug-delivery applications. TEM analysis also confirmed the spherical shape and small size (4-6nm) of P4-MET. Similarly, a PAMAM dendrimer loaded with rifampicin was characterized using the methods mentioned above to serve as a novel conjugate for targeting tuberculosis^20^.

The drug-release properties of P4-MET were carefully studied under varying pH conditions to simulate the physiological environment encountered by the nanocarrier across different tissues. The release of f-MET was fastest under acidic conditions (pH 4.5), characteristic of the tumor microenvironment, where pH is often lower due to metabolic activity and hypoxia. Under these acidic conditions, approximately 80% of f-MET was released within 50 hours, with near-complete release (100%) occurring after 100 hours. This rapid, pH-sensitive release is highly beneficial for cancer therapy, as it enables the rapid release of drugs at the tumor site while minimizing premature release elsewhere in the body. In contrast, under neutral pH (7.4), which resembles normal physiological conditions, the release of f-MET was significantly slower, with less than 20% released after 200 hours. This slow release at physiological pH ensures that P4-MET remains relatively stable in circulation, thereby reducing the risk of premature drug release before it reaches the tumor site. The pH-responsive release profile of P4-MET is particularly advantageous for targeting GBM cells, where the tumor’s acidic environment can trigger enhanced release of the encapsulated drug, ensuring targeted and effective therapeutic efficacy. These findings suggest that P4-MET can deliver f-MET in a controlled and efficient manner, providing an optimal drug-release profile for GBM treatment. The ability to selectively release the drug in the acidic tumor microenvironment could lead to higher concentrations of the therapeutic agent at the tumor site, potentially improving efficacy while reducing systemic toxicity. Likewise, Shi et al. scrutinize numerous studies demonstrating pH-dependent drug release from various polymers in tumor microenvironments (TMEs), which are relatively acidic compared to normal tissue (pH 7.4)^21^. In addition, TEM images revealed significant intracellular accumulation and alterations in mitochondrial morphology following f-Met and P4-MET treatments compared with control cells. Similarly, Bae et al. demonstrated ultrastructural alterations in HepG2 cells by TEM following PAMAM-O/Flag-apoptin treatment in HepG2 cells^22^.

Furthermore, proliferation inhibition assays demonstrated that P4-MET exhibited significantly lower IC50 values (40 to 250 times lower) compared to f-MET alone or a simple combination of f-MET and P4-OH dendrimers. This significant enhancement in anticancer activity is likely attributable to improved intracellular drug delivery facilitated by the P4-OH dendrimers. The dendrimer structure offers multiple points for drug conjugation, enabling efficient drug encapsulation and a more targeted release profile. Additionally, the smaller size of the P4-MET nanoparticles contributes to improved cellular uptake and greater efficacy, as they can more easily navigate the extracellular matrix and cell membranes. In accordance with our results, Kaveh Zenjanab et al. also demonstrated several-fold greater cytotoxicity of PAMAM dendrimer-conjugated Tirapazamine (TPZ) than that of free TPZ against breast cancer cells^23^. Moreover*, in vitro* studies were conducted to evaluate the cellular uptake of P4-MET in GBM cells. Cellular uptake studies revealed that P4-MET was efficiently internalized by cells, likely due to the enhanced permeability and retention (EPR) effect, which promotes nanoparticle accumulation at tumor sites. This enhanced uptake is a key advantage of using nanocarriers such as P4-MET, as it ensures that a higher proportion of the drug reaches tumor cells, a challenge often encountered with conventional drug delivery systems. The enhanced anti-cancer activity of P4-MET is indicative of its potential for treating GBM, a highly aggressive and infiltrating brain tumor. By improving the delivery of f-MET to target cells, P4-MET has the potential to overcome limitations of traditional drug formulations and deliver enhanced therapeutic outcomes for GBM patients. Similarly, many other studies have shown that the drug conjugated to a PAMAM dendrimer exhibits enhanced anticancer activity compared to its free form, due to greater uptake by cancer cells^24, 25^.

Additionally, metformin activates 5’-AMP-activated protein kinase (AMPK), which inhibits tumor cell growth and proliferation. Activated AMPK further inhibits AKT and mTORC1 phosphorylation and activation to restrain fatty acid ^26, 27^. Present results also showed time-dependent activation of AMPK expression levels due to a reduction in mitochondrial ATP and an elevation in Glycolytic ATP after both f-MET and P4-MET, as compared to control. These results are consistent with the earlier reports showing metformin induced energy stress due to the suppression of oxidative phosphorylation mediated generation of ATP in mitochondria, which further triggers the metabolic recompense via enhanced glycolytic flux^28, 29^.

Although it was fascinating to observe the opposite expression patterns of mTORC1 and its downstream effector, the ribosomal protein pS6, after f-MET and P4-MET treatment, this might be due to P4-MET’s more sustained inhibition of these pathways than f-MET. The inhibitory effect of this drug, mediated by AMPK activation, suggests that metformin could be used as a therapeutic agent in cancer treatment. Randomized controlled trials have shown improved survival in patients with pancreatic and lung cancer, demonstrating the clinical efficacy of metformin^30, 31^. Metformin can decrease mitochondrial-dependent oxygen consumption and ATP production, while enhancing glycolytic ATP production and lactate formation in glioma cells. Besides, mTOR pathway inhibition can also activate the REDD1 to induce cell death pathways in glioma cells ^4, 32, 33^. Our findings also showed a time-dependent upregulation of REDD1 following f-Met and P4-MET treatments. However, the upregulation was more pronounced after P4-MET treatment than with the free form, suggesting that the metformin nanoformulation has enhanced efficacy in targeting cancer cells.

Additionally, after identifying BOLA2B as a top differentially expressed gene following f-MET and P4-MET treatments in the RNA-seq analysis, we were intrigued to investigate correlations of BOLA2B with other signaling molecules, as BOLA2B was found to be highly expressed in malignant tumors compared with healthy tissue and could serve as a biomarker of poor prognosis across multiple cancers ^34^. Interestingly, the data showing that P4-MET downregulates p-mTORC1 levels without promoting BOLA2B, along with the positive correlations between mTORC1 and BOLA2B and GPX4, suggested ferroptosis-mediated GBM cell death after P4-MET treatment rather than canonical programmed cell death (apoptosis). Our data are in accordance with the present findings, which also showed induction of ferroptosis in GBM cells by inhibiting the AKT/mTORC1/GPX4 signaling pathway with fatostatin, which has antitumor effects across different tumor types ^35^. Recently, another study showed that mTORC1 inhibition combined with GPX4 inhibition is one of the most promising combinatorial approaches for mTOR-targeted anti-cancer therapies, and demonstrated that HDL-SCARB1-mediated suppression of ferroptosis is regulated by mTORC1 ^36^.

Many in vitro and in vivo murine models have been used to demonstrate the anti-cancer effects of metformin^37, 38^. However, to our knowledge, the present study is the first to demonstrate the antitumor effect of the metformin nanoformulation (P4-MET). After confirming the efficacy of P4-MET in glioma cells in vitro, we further evaluated its effect in the CT-2Aluc orthotopic mouse model. The in vivo safety profile of P4-MET was assessed by determining the maximum tolerated dose (MTD) in a mouse model. Mice were treated with P4-MET at a daily dose of 200 mg/kg, equivalent to the dose of f-MET alone, for 1 month. Notably, no signs of functional or histologic damage to vital organs were observed, indicating that P4-MET was well tolerated even at these high doses. Other studies have also confirmed that both metformin and dendrimer showed significant tolerance in vivo^39, 40^. In the present study, we used PAMAM dendrimers with OH surface groups, as they are preferred because they diffuse well through the BBB, do not adhere to surfaces, are hydrophilic, and are non-immunogenic^41^. The absence of toxicity suggests that the dendrimer-based nanocarrier does not induce significant adverse effects at therapeutic doses, a crucial prerequisite for the clinical translation of P4-MET. The favorable safety profile of P4-MET is a promising indicator of its potential use in clinical settings, as it demonstrates that the formulation can be administered over extended periods without causing significant harm. High dose tolerance is critical in cancer therapy, where higher doses are often required to achieve therapeutic efficacy. This aspect of P4-MET further underscores its suitability as a safe and effective drug delivery system for cancer treatment.

The therapeutic efficacy of P4-MET was evaluated in an orthotopic syngeneic GBM mouse model. Survival studies showed that treatment with P4-MET significantly prolonged overall survival (OS) in mice compared with f-MET alone. The median OS of the P4-MET-treated group was substantially longer, demonstrating the enhanced therapeutic potential of P4-MET. This improvement in survival is likely due to enhanced anticancer activity and targeted drug delivery achieved by the PAMAM dendrimer-based system. Moreover, when P4-MET was combined with radiation therapy (RT), the mice exhibited an even more pronounced increase in OS. Two-thirds of the mice treated with the combination of P4-MET and RT achieved tumor-free survival for over two months. The combination of P4-MET with RT may enhance the therapeutic outcome by increasing the effectiveness of both treatments. RT is known to induce DNA damage and cell death in tumor cells. When combined with P4-MET, it may enhance the drug’s efficacy by improving its penetration into the tumor and targeting cancer cells more effectively. Metformin reduced tumor growth and mitigated drug resistance in preclinical studies.

Additionally, numerous clinical trials have reported that metformin may improve overall survival when combined with other therapies. In line with our findings, Sesen et al. demonstrated that metformin delays tumor growth in nude mice in vivo and acts as a chemo- and/or radiosensitizer. The present results suggest that combining P4-MET with RT holds excellent promise as a treatment strategy for GBM, offering a novel and effective therapeutic approach to improve patient outcomes in this challenging cancer. There is a substantial body of evidence showing that combining radiotherapy or chemotherapeutic agents with drugs targeting cellular metabolism is a desirable therapeutic alternative for cancer. To date, there have been no other reports of combining radiation therapy with a nano-formulation of metformin targeting GBM, as shown in the present findings.

Furthermore, Pharmacokinetics and drug distribution studies were performed using fluorescence-tagged P4-MET to trace its biodistribution *in vivo*. P4-MET was widely distributed in all vital organs, with the highest accumulation observed in the kidneys and liver. These organs play a significant role in the metabolism and excretion of nanoparticles, consistent with their functions in the body. As reported in many studies, PAMAM-OH can penetrate the BBB^46, 47^; notably, P4-MET can also cross the blood-brain barrier (BBB), an essential feature for treating brain tumors such as GBM. The ability of P4-MET to cross the BBB is a significant advantage, as it enables direct delivery of therapeutic agents to the brain tumor site. Interestingly, the combination of P4-MET with radiation therapy (RT) resulted in a significant increase in drug distribution in the brain, with fluorescence signals nearly 3-fold higher after RT. This enhanced drug penetration into the brain following RT suggests that BBB permeability may increase after radiation, allowing more efficient drug delivery to the tumor site. This finding further supports the potential of combining P4-MET with RT to improve therapeutic outcomes in GBM treatment. These pharmacokinetic findings are crucial for the development of P4-MET as a therapeutic agent for GBM.

Moreover, many recent studies have highlighted metformin’s effects on T cells in the tumor microenvironment (TME) and shown that it reduces hypoxia-induced immunosuppression^48–50^. Our IHC results of mouse brain sections stained with CD8a and CD4 also showed significant upregulation of T-cells after P4-MET treatment compared to f-MET. Additionally, metformin is known to polarize M2-like (anti-inflammatory) macrophages toward M1-like (pro-inflammatory) macrophages^51, 52^. Our IHC findings also showed a significant decrease in CD206-positive cells after P4-MET treatment, a specific marker of the M2 Phenotype of macrophages, thereby indicating its anti-inflammatory effect. To recapitulate, these results suggest that the anticancer effects of metformin and P4-MET may be partly mediated by the host immune system.

After 4 months of initial P4-MET and brain RT treatment, 4 mice survived and were included in our subsequent rechallenge study, in which we reimplanted CT-2Aluc cells. It was interesting to find that only 1 of 4 mice developed tumors, whereas all naive mice in the control group developed tumors (as determined by IVIS imaging). Results indicate the formation of long-term immune memory after P4-MET+RT treatment, primarily involving T_EM of CD4 and CD8 in both blood and spleen samples from rechallenged mice compared with control mice. Metformin demonstrated protection against the apoptosis of CD8+ tumor-infiltrating lymphocytes (TILs). It restored the exhausted PD-1-Tim-3+ CD8+ TILs by shifting them from a central memory (CM) phenotype to an effector memory (T_EM) phenotype. Consistent with our results, Eikawa et al. also demonstrated that metformin leads to T_EM dominance over TCM in the MO5 model. These significant alterations in the properties of CD8+ TILs are thus correlated with metformin’s anti-tumor mechanism^48^. Our results also confirmed that T_EM is more accountable for tumor growth inhibition than CM after P4-MET and RT treatment. The increase in T_EM cells indicates a shift toward a more robust and effective immune response, which could be critical for combating glioblastoma recurrence. Conversely, we hypothesized that metformin would have a minimal impact on cold tumors, such as GBM. Thus, combination therapy with P4-MET, RT, and/or other immune checkpoint inhibitors is considered a promising approach for targeting cold tumors such as GBM.

These findings underscore the potential of P4-MET not only as a direct therapeutic agent for glioblastoma but also as a strategy to enhance immune-mediated tumor elimination by modulating the host immune system. To recapitulate, the present study demonstrated that P4-MET can cross the BBB, distribute effectively at the tumor site, and enhance radiosensitization of GBM with RT, indicating that P4-MET is a promising candidate for targeting brain tumors, specifically GBM.

## Conclusion

P4-MET exhibited a favorable toxicity profile compared to f-MET at equivalent doses while demonstrating significantly enhanced anti-tumor efficacy in both in vitro and in vivo GBM models. The conjugation of metformin to P4-OH dendrimers enhanced penetration across the blood-brain barrier, resulting in greater drug accumulation in the tumor microenvironment. Additionally, combining P4-MET with radiotherapy further enhanced brain penetration and therapeutic efficacy, resulting in prolonged overall survival in glioblastoma-bearing mice, depicting radio-sensitizing efficacy. Additionally, this drug has the potential to simultaneously downregulate tumor-supporting immune cells and upregulate tumor-targeting immune cells. These findings highlight P4-MET as a promising nanotherapeutic for GBM treatment, offering the dual advantages of direct tumor suppression and indirect immune modulation while minimizing systemic toxicity. Future studies will optimize the formulation and evaluate its translational potential in preclinical and clinical settings.

## Acknowledgement

The authors would like to thank Deepti Negi and Nicholas Conoan of the Electron Microscopy Core Facility (EMCF) at the University of Nebraska Medical Center for technical assistance. The EMCF is supported by state funds from the Nebraska Research Initiative (NRI) and the University of Nebraska Foundation, and institutionally by the Office of the Vice Chancellor for Research.

The UNMC Flow Cytometry Research Facility is administered through the Office of the Vice Chancellor for Research and supported by state funds from the Nebraska Research Initiative and The Fred and Pamela Buffett Cancer Center’s National Cancer Institute Cancer Support Grant (P30 CA036727). Major instrumentation has been provided by the Office of the Vice Chancellor for Research, The University of Nebraska Foundation, the Nebraska Bankers’ Fund, and the NIH-NCRR Shared Instrument Program.

## Author Contributions

All authors contributed to conception and design. S.M., F.W., T.K.B., and S.R. conducted experiments and analysis. G.M.C. performed all the bioinformatics analysis of the RNA-Seq data. S.M., F.W., T.K.B., S.R., K.B., Y.J., and C.K.Z. provided a critical interpretation. S.M., S.R., K.B., Y.J., and C.K.Z. wrote the manuscript with input from all authors. All authors reviewed and approved the final manuscript.

## Ethics Approval

All mouse experiments/procedures were duly approved and performed as per the Institutional Animal Care and Use Committee (IACUC) guidelines in accordance with the National Institute of Health (NIH). The animals were maintained and operated in accordance with IACUC standards at the University of Nebraska Medical Center (UNMC) animal facility. The protocol number was 22-074-07-FC.

## Funding

The project was supported by the National Institute of General Medical Sciences (1U54GM115458-01) to CK.Z. and the Robert Holmes Foundation to C.L.

## Conflict Of Interest

The authors declare no competing financial interests.

## Data Availability

All data supporting this study are available from the corresponding author upon reasonable request.

## Notes

### Competing Interest Statement

The authors have declared no competing interest.

### Summary of Updates

There was an error in the corresponding author's name. we corrected that only

