## Supplementary Figure 1-5 for "Enhanced Radio-sensitization of Glioblastoma using a Dendrimer-Based Metformin Nano-formulation through Direct Tumor Suppression and Indirect Immune Modulation"

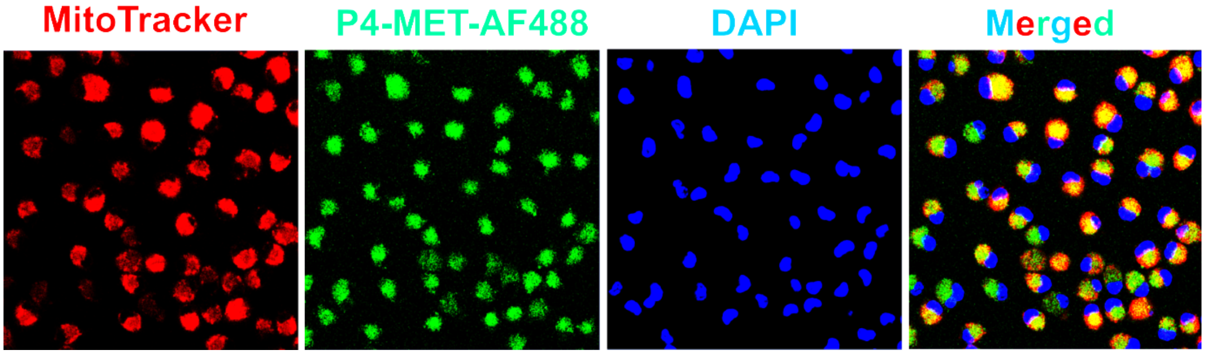


**Figure S1.** The intracellular localization and distribution of the P4-MET tagged with AF488 by bone marrow derived macrophages (BMDM) after 24 hours treatment was evaluated by confocal microscopy. Mitochondria were stained with MitoTracker Red for 30 minutes followed by the nuclear counterstain with DAPI.

**Supplementary Figure 2**

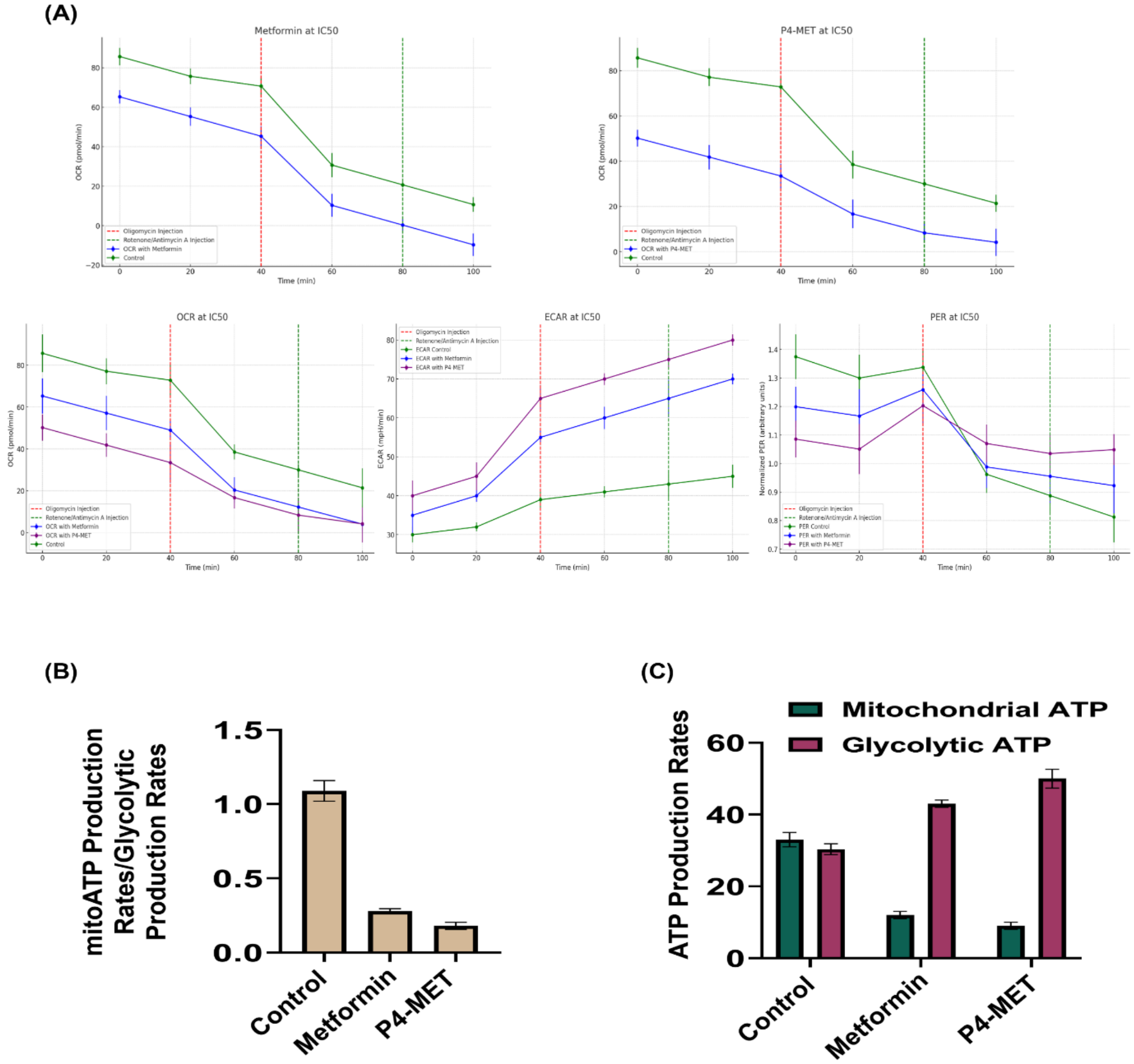


**Figure S2. (A)** Seahorse XFe96/XF Pro microplates were used to evaluate mitochondrial and glycolytic ATP production through Agilent analyzer. **(B)** The results show a remarkable reduction in the ratio of mitochondrial ATP production rates and glycolytic ATP production rates. **(C)** The ATP production rate showed decline in mitochondrial ATP and elevation in glycolytic ATP production after f-MET and P4-MET treatment as compared to control. These data demonstrate the robust measure of metabolic reprogramming induced by both f-Met and P4-MET.

**Supplementary Figure 3**

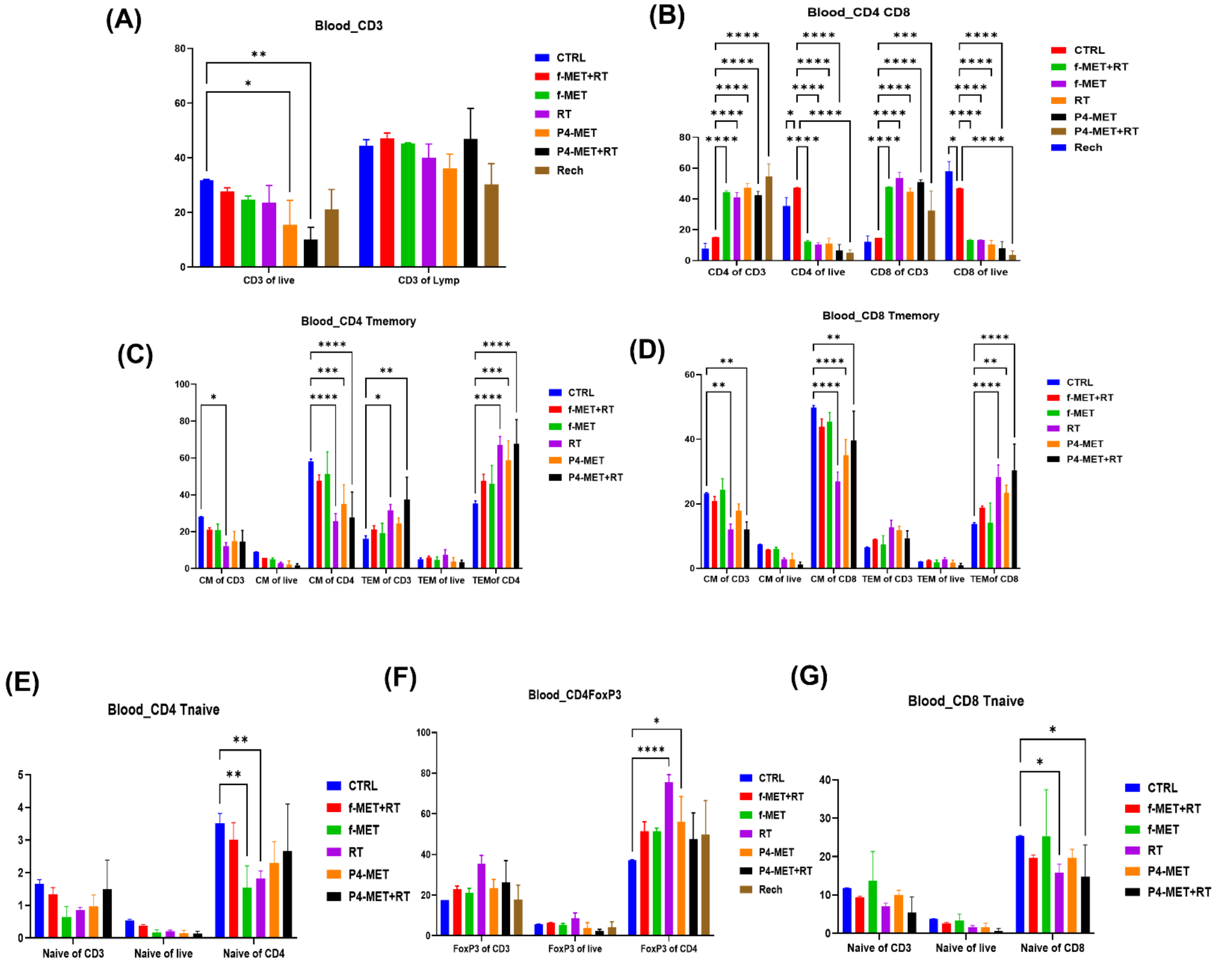


**Figure S3.** Flow cytometry analysis of circulating immune cell populations in peripheral blood samples across different treatment groups, including control (untreated), f-MET, f-MET+RT, RT, P4-MET, P4-MET+RT, and rechallenged (Rech). (A)Percentage of CD3+ T cells out of total live cells and total lymphocytes. (B) Proportions of CD4+ and CD8+ T cell subsets relative to CD3+ or live cells. (C) Proportions of central memory (CM) and effector memory (TEM) populations within CD4+ T cells. (D) Quantification of central memory and effector memory subsets within CD8+ T cells. (E) Percentage of naïve CD4+T cells within CD3+, live, and CD4+ cells. (F) Levels of circulating FOXP3+ regulatory T cells (Tregs) among CD3+, live, and CD4+ populations. (G) Percentage of naïve CD8+T cells within CD3+, live, and CD8+ populations.

**Supplementary Figure 4**

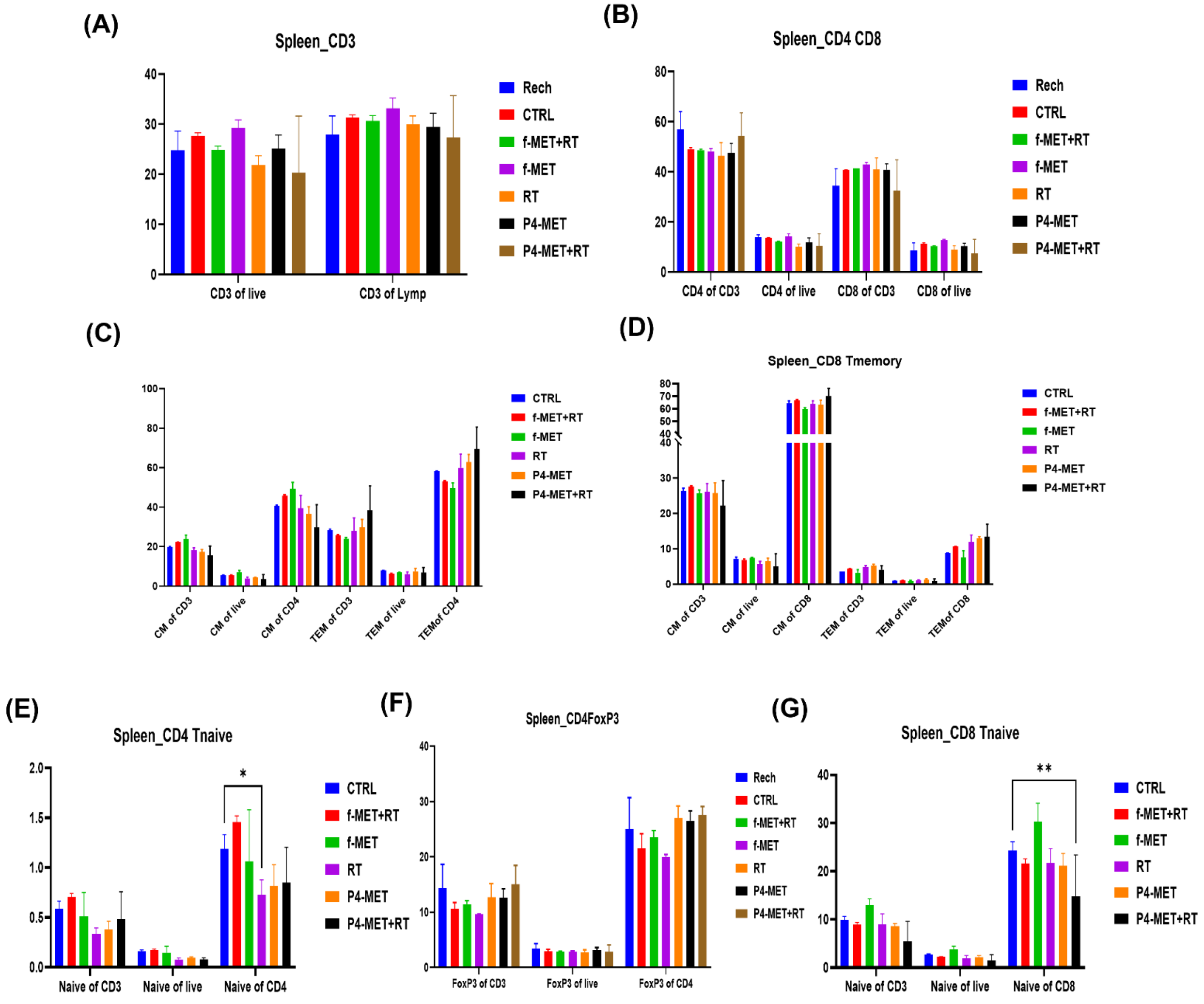


**Figure S4.** Flow cytometry analysis of immune cell populations in the spleen samples across different treatment groups, including rechallenged (Rech), control (untreated), f-MET, f-MET+RT, RT, P4-MET, and P4-MET+RT. (A)Percentage of CD3+ T cells out of total live cells and total lymphocytes. (B) Proportions of CD4+ and CD8+ T cell subsets relative to CD3+ or live cells. (C) Proportions of central memory (CM) and effector memory (TEM) populations within CD4+ T cells. (D) distribution of central memory and effector memory subsets within CD8+ T cells. (E) Percentage of naïve CD4+T cells within CD3+, live, and CD4+ cells. (F) Levels of FOXP3+ regulatory T cells (Tregs) among CD3+, live, and CD4+ populations. (G) Percentage of naïve CD8+T cells within CD3+, live, and CD8+ populations.

### Supplementary Figure 5

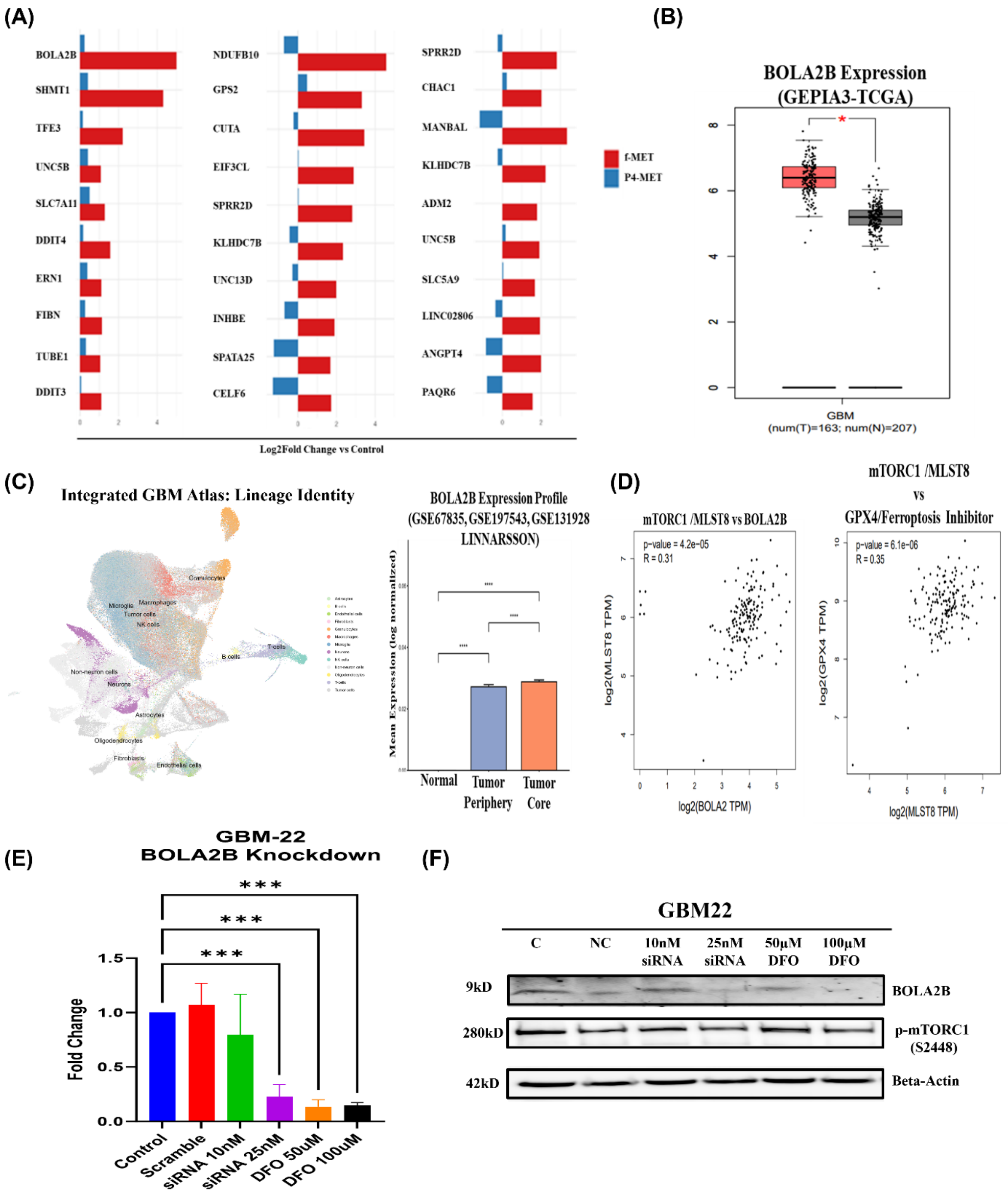

**Figure S5.** Differential gene expression (DEG) analysis in GBM cells after f-Met and P4-MET treatment at different time points. (**A**) RNA-seq analyses identified BOLA2B as the top differentially expressed gene between the treatment groups at four hours. (**B**) TCGA data showed significant overexpression of BOLA2B in tumors compared with normal brain tissue. (**C**) Single-cell RNA-seq databases from GBM patients also confirmed BOLA2B overexpression in both the periphery and core of GBM relative to the healthy brain. (**D**) Correlation analysis between BOLA2B and mTORC1, and between mTORC1 and GPX4 (Ferroptosis inhibitor), revealed positive correlations (p-values $< 0.0001$). (**E**) RT-qPCR results confirmed successful knockdown of BOLA2B expression by BOLA2B-specific siRNAs compared with the control (untreated) and negative control (scrambled siRNA). (**F**) Immunoblotting further showed significant downregulation of BOLA2B protein expression by 25nM siRNA and 100 μM DFO treatment, respectively, along with reduced phosphorylation of mTORC1 (S2448).
